# Analysis of regulatory element evolution between human and mouse reveals a lack of *cis-trans* compensation

**DOI:** 10.1101/847491

**Authors:** Kaia Mattioli, Winona Oliveros, Chiara Gerhardinger, Daniel Andergassen, Philipp G. Maass, John L. Rinn, Marta Melé

## Abstract

Gene expression differences between species are driven by both *cis* and *trans* effects. Whereas *cis* effects are caused by genetic variants in close proximity to the target gene, *trans* effects are due to distal genetic variants that affect diffusible elements such as transcription factors. Previous studies have mostly assessed the impact of *cis* and *trans* effects at the gene level. However, how *cis* and *trans* effects differentially impact regulatory elements such as enhancers and promoters remains poorly understood. Here, we used massively parallel reporter assays to directly measure *cis* and *trans* effects between human and mouse embryonic stem cells at thousands of individual regulatory elements. Our approach revealed that *cis* effects are widespread across regulatory elements, and the strongest *cis* effects are associated with the disruption of motifs recognized by strong transcriptional activators. Conversely, we found that *trans* effects are rare but stronger in enhancers than promoters, and can be attributed to a subset of transcription factors that are differentially expressed between human and mouse. While previous studies have found extensive co-occurrence of *cis* and *trans* effects in opposite directions that stabilize gene expression throughout evolution, we find that *cis-trans* compensation is uncommon within individual regulatory elements. Thus, our results are consistent with a model wherein compensatory *cis-trans* effects at the gene level are explained by *cis* and *trans* effects that separately impact several regulatory elements rather than *cis-trans* effects that occur simultaneously within a single regulatory element. Together, these results indicate that studying the evolution of individual regulatory elements is pivotal to understand the tempo and mode of gene expression evolution.

## INTRODUCTION

Since King and Wilson suggested that changes in transcriptional regulation underlie phenotypic differences between species^1^, it has become clear that changes in gene expression are heritable and often play a role in the evolution of phenotypes^2,3^. Changes in non-coding regulatory elements—including promoters and enhancers—are particularly important in driving the evolution of gene expression^4,5^. Two primary mechanisms are responsible for the evolution of gene expression: *cis* effects and *trans* effects. *Cis* effects are due to genetic variants that are in linkage disequilibrium with the target gene; for example, genetic variants located in gene promoters or enhancers that affect transcription factor (TF) binding sites. Conversely, *trans* effects are driven by diffusible elements (such as TFs) that are distal and unlinked to the genes they affect. Any given gene can be subject to *cis* effects, *trans* effects, or both^6^.

Much work has assessed the contribution of *cis* and *trans* effects on the evolution of gene expression. One of the most common approaches has been to perform allele-specific RNA sequencing of F1 hybrid offspring, which can separate regulatory variants acting in *cis* (which show allele-specific effects in the hybrid) from those acting in *trans* (which affect both hybrid alleles equally)^6^. These studies have assessed both intra- and inter-species variation in gene expression across a variety of taxa, including yeast^7,8^, insects^9,10^, plants^11^, and mice^12^. These and other studies have shown a predominance of *cis* effects, but highlighted a role for *trans* effects that varied across taxa. Moreover, *cis* and *trans* effects were found to often occur simultaneously and affect target gene expression in opposite directions^12^. This so-called “compensation” between *cis* and *trans* effects is thought to stabilize gene expression throughout evolution^6^. A major limitation of these studies, however, is that while they can assign *cis* and *trans* effects to target genes, they cannot disentangle effects at individual regulatory elements. Studies on regulatory element evolution have found that the number of regulatory elements—especially enhancers—that target a gene influence the tempo and mode of gene expression evolution^5,13^. However, only small scale studies have examined how *cis* and *trans* effects drive differences in regulatory element activities across species^14,15^.

The development of massively parallel reporter assays (MPRAs) has revolutionized our ability to dissect the regulatory element code^16,17^. Indeed, MPRAs have been used to measure regulatory element activity of thousands of sequences across tissues^18^, species^15^, and allelic variants^18–20^. In this work, we use MPRAs to quantitatively investigate *cis* and *trans* effects across thousands of individual regulatory elements including enhancers, promoters of protein-coding genes, and promoters of long non-coding RNA (lncRNA) genes. We perform MPRAs in similar cellular environments from two mammalian species—embryonic stem cells (ESCs) from human and mouse—to perform a systematic analysis of *cis* and *trans* effects at thousands of individual regulatory elements simultaneously.

## RESULTS

### Designing an MPRA to measure regulatory element evolution

To investigate regulatory element evolution between human and mouse, we first defined regulatory elements in both species using a set of robust transcription start sites (TSSs) defined by the FANTOM5 consortium^21^. We categorized these TSSs into three major biotypes: (1) eRNAs (RNAs emerging from bidirectionally transcribed enhancers that do not overlap protein-coding genes), (2) lncRNA promoters, and (3) mRNA promoters (see methods). We then projected these TSSs onto the genome of the other species (i.e., human TSSs were projected onto the mouse genome and vice versa). We classified TSSs as “sequence orthologs” if we were able to reciprocally map the TSS between the two species. We further classified the “sequence ortholog” TSSs as conserved TSSs if the aligned region in the other species (+/−50 bp from the TSS) contained evidence of an active TSS (**Figure 1A**; see methods). As expected, the proportion of TSS that were sequence orthologs and conserved were both highest in mRNAs and lowest in eRNAs (**Figure 1B; Supplemental Figure S1**). Despite moderate levels of sequence orthology in eRNAs and lncRNAs, both biotypes exhibited very high activity turnover, with only 7% and 31% of human eRNA TSSs and lncRNA TSSs being conserved in mouse, respectively.

**Figure 1:**
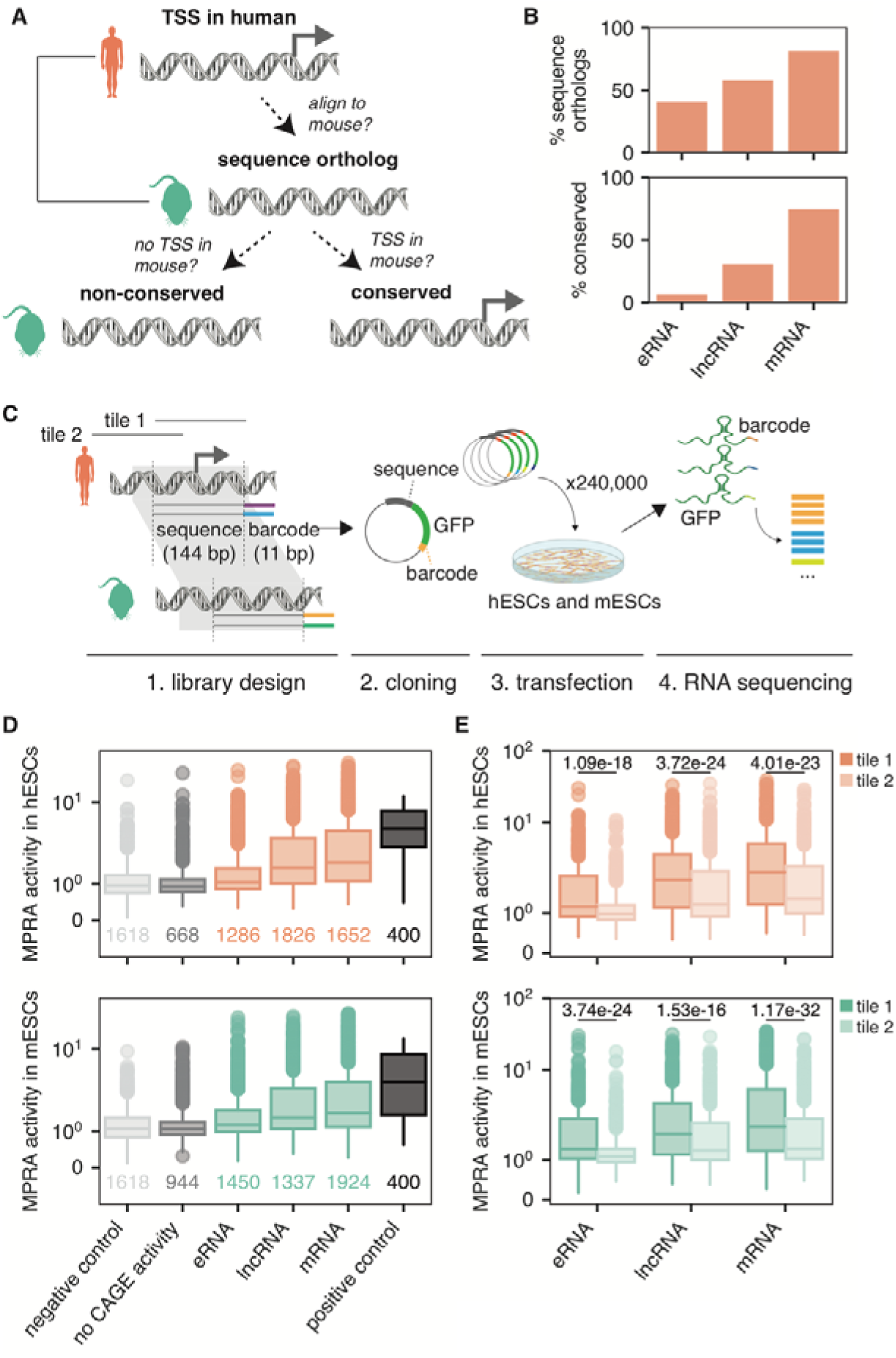
MPRA in human and mouse ESCs recapitulates endogenous gene expression patterns. **A:** Schematic depicting the definitions of a sequence ortholog and conserved/non-conserved TSSs. Sequence orthologs are TSS regions whose sequences can be reciprocally mapped to the other species. Conserved TSSs are a subset of sequence orthologs that also overlap a TSS in the other species (defined as having ≥ 10 CAGE reads in ≥ 1 sample), whereas non-conserved TSSs do not. **B:** Percentage of human-to-mouse sequence orthologs and conserved TSSs broken up by biotype. **C:** Schematic of MPRA design. Tile 1 overlaps the assigned TSS (114 bp upstream to 30 bp downstream) whereas tile 2 does not (228 bp upstream to 84 bp upstream). **D:** MPRA activities of human (top) and mouse (bottom) TSSs in their native contexts, human ESCs and mouse ESCs, respectively, broken up by whether they have endogenous CAGE activity and then by biotype and compared to negative controls (random sequences) and positive controls (CMV promoter regions). **E:** MPRA activities of TSS-overlapping tile (tile 1) compared to upstream tile (tile 2) across all human biotypes (top) and mouse biotypes (bottom). P-value shown is from a two-sided Mann Whitney test.

To systematically assess the contribution of *cis* and *trans* effects to the evolution of thousands of regulatory elements simultaneously, we performed a massively parallel reporter assay (MPRA) (**Figure 1C**). MPRAs measure the transcriptional activities of designed sequences in a cell type of interest, and thus enable us to test both *cis* effects (how do orthologous sequences compare within a given cellular environment) and *trans* effects (how do different cellular environments affect a given sequence). As early development is known to play a key role in evolutionary processes^22^, we chose to perform the MPRA in a developmentally-relevant cell type: human and mouse ESCs. Thus, we selected 3,327 pairs of orthologous regulatory elements between human and mouse, all of which had endogenous activity in either human or mouse ESCs or both (**Supplemental Figure S2**; **Supplemental Table S1**; see methods). The full list of regulatory elements in our library can be found in **Supplemental Table S2**. To ensure that we covered all regulatory activity sequence surrounding the TSS, we designed two oligonucleotide tiles for each TSS (**Figure 1C**). All told, our library included 13,533 sequences to test (**Supplemental Table S3**). To control for technical variation across sequencing measurements, each element was represented a minimum of 13 times, each time with a different barcode. We also included randomly-generated sequences as negative controls (with 3 barcodes each) as well as tiled regions of the cytomegalovirus (CMV) promoter as positive controls (with 60 barcodes each), resulting in a final library of 181,065 unique oligonucleotides (**Supplemental Table S4**). We performed three biological replicates each in human ESCs (hESC) and mouse ESCs (mESCs) and confirmed that replicates of hESCs and mESCs clustered separately (**Supplemental Figure S3** and **Supplemental Figure S4**). We then removed barcodes with low counts, resulting in a set of 2,952 regulatory sequence pairs that were well represented in our data (see methods).

We next quantified each sequence’s ability to drive transcription in the MPRA experiment—termed “MPRA activity”—using MPRAnalyze^23^. Briefly, MPRAnalyze uses a graphical model to estimate the rate of transcription of each sequence in the library by comparing RNA counts for each barcode to input DNA counts for each barcode. To determine whether our MPRA was able to recapitulate biological signal, we compared the MPRA activity of each regulatory element in its native context (human sequences in hESCs and mouse sequences in mESCs) to negative and positive control sequences (see methods). TSSs were more active than negative controls and mimicked endogenous activity levels: eRNAs had the lowest activity while mRNAs had the highest activity (**Figure 1D**).

We then compared the activity of the annotated TSS-overlapping tiles (tile 1) to the upstream tiles (tile 2) (**Figure 1C**). As expected, across all biotypes, annotated TSS-overlapping tiles were significantly more active in their native context than the upstream tiles (**Figure 1E**). In 18% of regulatory element pairs, however, the upstream tile was more active than the TSS-overlapping tile in both species’ (**Supplemental Figure S5**), likely due to slight misannotation of the exact TSS location. Thus, while FANTOM5-defined TSSs are highly accurate, including additional upstream regions in the MPRA can help to refine core promoter locations. We therefore assigned each of the 2,952 regulatory element pairs a single representative tile to use in both species: we always used the annotated TSS-overlapping tile except in those cases where the upstream tile had more activity in both species. Among those, 1,644 pairs (55%) had significant MPRA activity (MPRA q-value < 0.05) in at least 1 native context. We limited all of our subsequent analyses to this set of 1,644 active sequence pairs (3,288 sequences total).

### *Cis* effects are common and associated with evolutionary turnover

Differences in regulatory element activity between species could be due to differences in DNA sequence (*cis* effects) or cellular context differences (*trans* effects) between the species or both. We decided to focus first on *cis* effects, which can be attributed to differences in DNA sequence alone. We defined *cis* effects as the MPRA activity differences between orthologous sequence pairs in the same cellular environment (**Figure 2A**). To calculate *cis* effects, we used MPRAnalyze to test for MPRA activity differences between pairs of orthologous regulatory elements. An advantage of using MPRAnalyze is that it is able to use information from null differential controls to inform its comparative model. The ideal null differential controls are pairs of identical sequences tagged with different barcodes. We therefore leveraged our CMV tiles, each of which was attached to 60 barcodes, to create our null differential controls by down-sampling barcodes (**Supplemental Figure S6** and **Supplemental Figure S7**; see methods). As expected, orthologous regulatory element pairs had higher *cis* effect sizes than null differential controls in both hESCs and mESCs (**Figure 2B**). Overall, 40% of the 1,644 tested regulatory element pairs showed a significant *cis* effect in hESCs, mESCs, or both (empirical FDR < 0.1) (**Figure 2C**; see methods).

**Figure 2:**
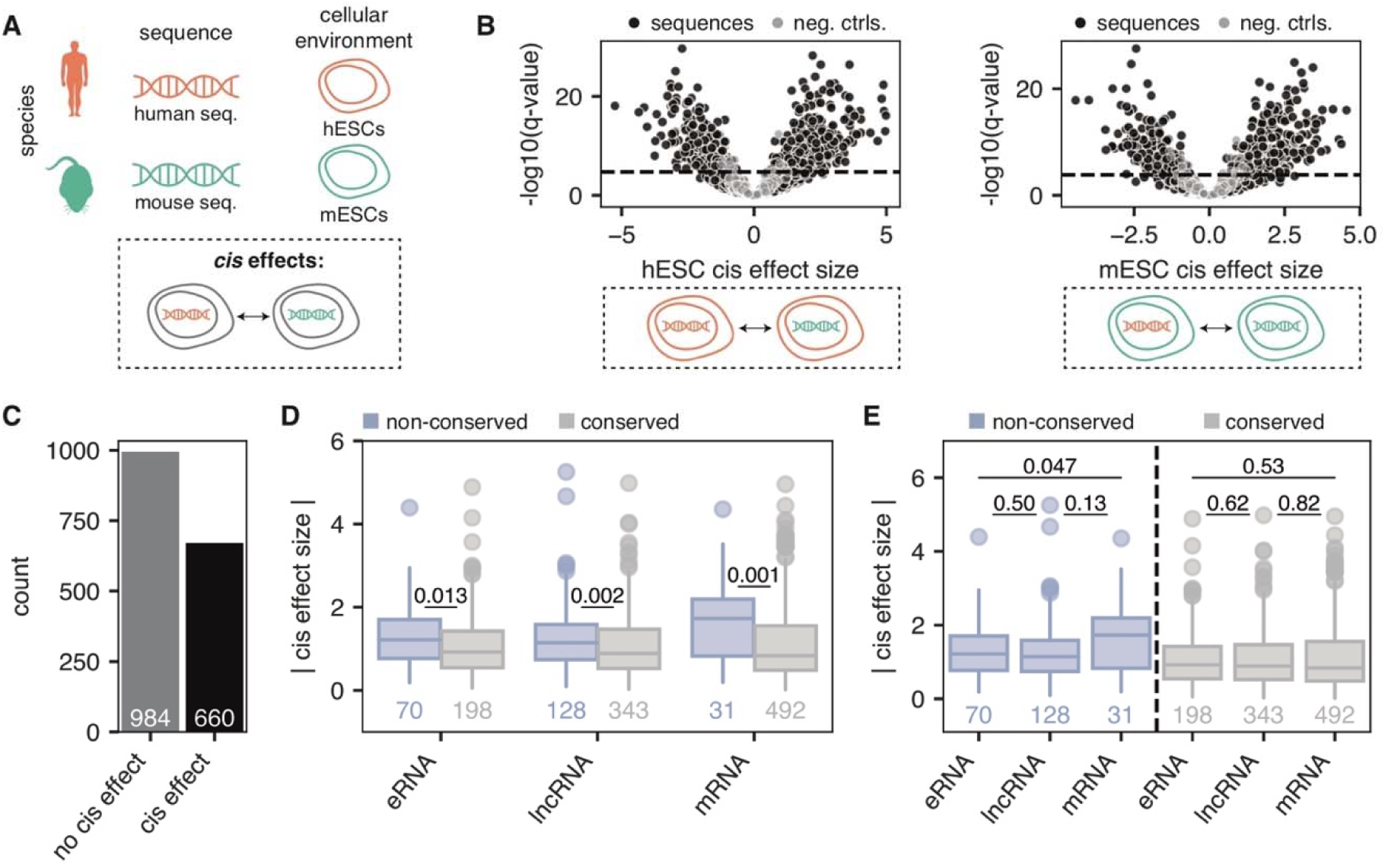
40% of orthologous regulatory elements show significant *cis* effects. **A:** Schematic depicting the definition of a *cis* effect: MPRA activity differences between human sequences and mouse sequences while keeping the cellular environment constant. **B:** Volcano plot showing the cis effect sizes (log2 foldchanges in activity between sequences in hESCs, left and mESCs, right) of orthologous sequences (black) and null differential controls (gray). Horizontal line depicts an empirical FDR cut-off of 0.1, calculated using null differential controls (see methods). **C:** Count of orthologous sequence pairs with significant *cis* effects. **D:** Absolute cis effect sizes across biotypes, broken up into non-conserved TSSs (blue) and conserved TSSs (gray). P-values shown are from a two-sided Mann Whitney test. **E:** Same as (D), but this time comparing differences across biotypes. P-values shown are from a two-sided Mann Whitney test.

We next sought to examine how *cis* effects differ across biotypes, including conserved and non-conserved TSSs. Within each biotype, non-conserved regulatory element pairs showed significantly higher *cis* effects than conserved regulatory element pairs (**Figure 2D**). We confirmed that our non-conserved pairs are *bona fide* non-conserved regulatory elements, as the non-conserved pairs we had defined (non-conserved TSSs and their orthologous sequence in the other species that lacked a TSS) had higher pairwise alignment scores than they did to the closest TSS in the other species (**Supplemental Figure S8**). Thus, these non-conserved regulatory elements are not due to misalignments between genomes. *Cis* effect sizes across biotypes were relatively uniform (**Figure 2E**). However, non-conserved mRNA TSSs showed the highest *cis* effect sizes (**Figure 2E**), consistent with the idea that the largest jump in activity is from mRNA TSSs—which have the highest activity out of all biotypes—to sequences without a TSS at all.

### *Cis* effects are associated with disruption of certain TF motifs

*Cis* effects are often caused by disruption of motifs that are recognized by sequence-specific transcription factors (TFs). Thus, we next sought to determine the relationship between *cis* effects and TF motifs. Previous work showed that only a subset of TF motifs can be reliably associated with MPRA activity variance^24^. Thus, we selected a set of 466 motifs from TFs that are expressed in hESCs and mESCs and are associated with MPRA activity (**Supplemental Figure S9**; **Supplemental Table S5**) either as activators or as repressors for further analysis. As expected, regulatory element pairs showing no *cis* effects shared more TF motifs than sequence pairs with significant *cis* effects (**Figure 3A**), reinforcing the notion that the more TF motifs two sequences have in common, the more similar their activity levels. In addition, we found 17 individual motifs were significantly associated with *cis* effects when disrupted. The majority of these motifs were predicted activators and enriched in mRNAs (**Figure 3B**); indeed, several of the strongest effect sizes could be attributed to the ETS transcription factors, including the oncogenic TF ETV1^25^ (**Figure 3C**). However, we also found a subset of motifs that were predicted repressors and enriched in eRNAs (**Supplemental Figure S10**). Thus, while *cis* effects can generally be attributed to the disruption of strong activating motifs, in rarer cases, *cis* effects are due to the disruption of weak repressive motifs. While this may reflect real biological effects, it may also be due to the fact that MPRAs are more powered to detect activators over repressors^24,26^.

**Figure 3:**
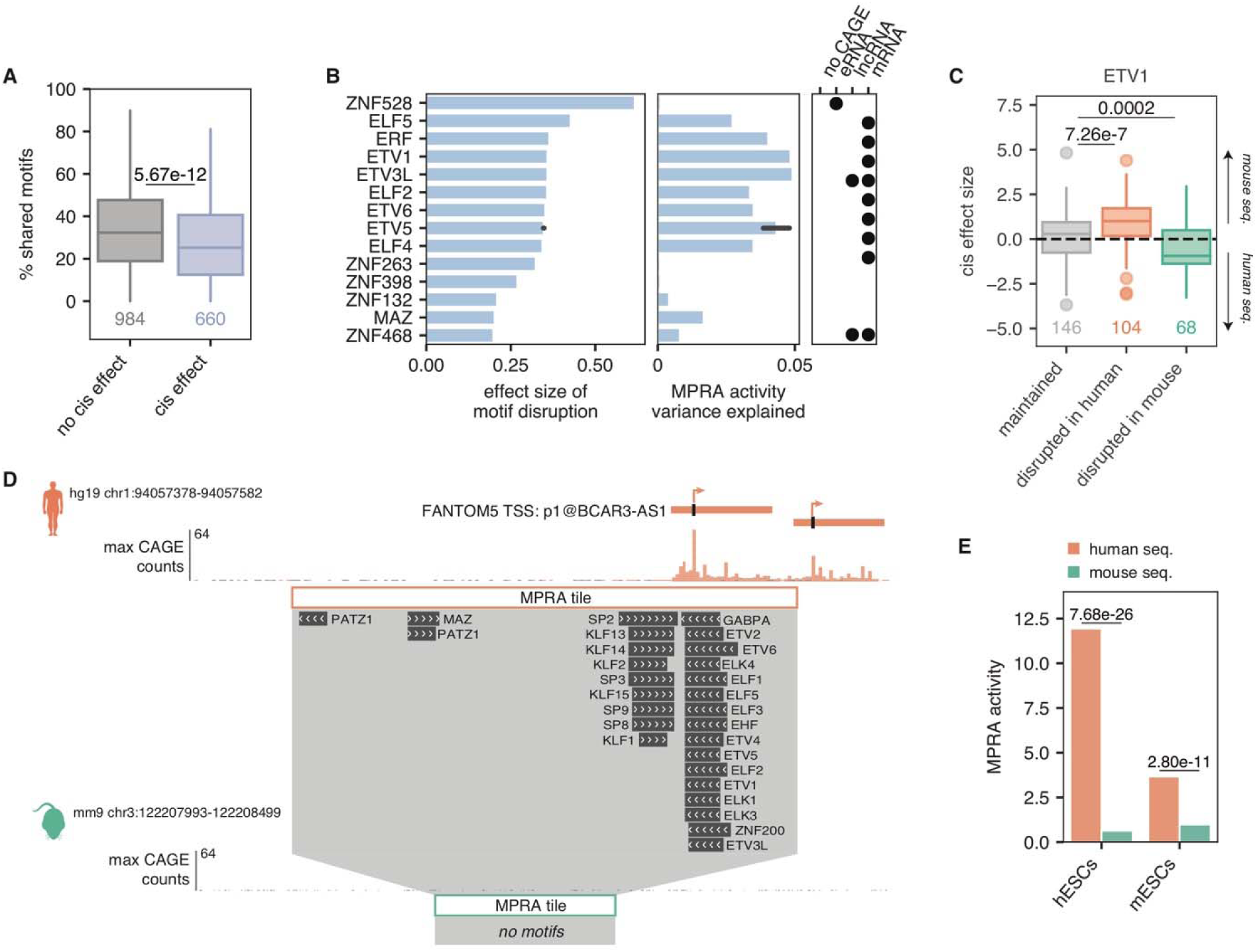
Disruption of certain motifs is associated with *cis* effects. **A:** Percentage of shared motifs in tiles that show cis effects vs. those that do not. P-value shown is from a two-sided Mann Whitney test. **B:** Plot showing the activating motifs whose disruption is significantly associated with *cis* effects (FDR < 0.05). Left: effect size associated with motif disruption. Middle: additional variance in MPRA activity explained by the TF. Right: enrichment of a given TF motif across biotypes, as determined by a Hypergeometric test. Black dots denote significant enrichment (FDR < 0.05). The ETV5 TF has two “best” motifs according to the curated Lambert et al. TF list and therefore the average of these two motifs are plotted, with the bootstrapped 95% confidence interval shown. **C:** Relationship between *cis* effect sizes and the ETV1 motif, where “maintained” are sequence pairs that both have the ETV1 motif, “disrupted in human” are pairs where the ETV1 motif is present in mouse but not in human, and “disrupted in mouse” are pairs where the ETV1 is present in human but not in mouse. A *cis* effect size > 0 indicates the mouse sequence has higher activity whereas a *cis* effect size < 0 indicates the human sequence has higher activity. P-values shown are from a two-sided Mann Whitney test. **D:** Genome browser screenshot of an example locus showing a *cis* effect. Only motifs that were found to explain ≥1% of the variance in MPRA activity are shown. **E:** MPRA activities for human sequence (orange) and mouse sequence (green) in hESCs and mESCs for the locus shown in D. P-values shown are the q-values calculated by MPRAnalyze.

An example of a *cis* effect can be seen at the TSS for the human-specific lncRNA BCAR3-AS1. In human, the strongest core promoter region of BCAR3-AS1 contains many strong activating motifs, including several ETS motifs (**Figure 3D**). The orthologous region in the mouse, however, shows no CAGE activity and lacks these motifs, as there is no orthologous lncRNA in the mouse (**Figure 3D**). As expected, the pairwise alignment score between the human BCAR3-AS1 TSS and region and the region shown in **Figure 3D** is higher than the alignment score to the nearest mouse TSS (160.9 compared to 138.5), indicating that our MPRA tiles are correctly aligned. In our MPRA, this pair shows a significant *cis* effect: the human sequence is significantly more active than the orthologous mouse sequence in both hESCs and mESCs (**Figure 3E**). Collectively, our results show that *cis* effects are common—especially in regulatory element pairs that show activity changes between species—and associated with disruption of specific TF motifs.

### *Trans* effects are rare and highest in eRNAs

After quantifying *cis* effects, we next sought to quantify *trans* effects. We defined *trans* effects as the difference in MPRA activity driven by differences in cellular environment alone and measured them by quantifying MPRA activity differences between hESCs and mESCs while keeping the sequence constant (**Figure 4A**). As with *cis* effects, human and mouse regulatory elements showed higher *trans* effects than null differential controls (**Figure 4B**). Overall, 18% of the 1,617 filtered regulatory element pairs with significant activity showed a significant *trans* effect in the human sequence, the mouse sequence, or both (**Figure 4C**). Compared to *cis* effect sizes, however, *trans* effect sizes were much lower. In addition, unlike *cis* effects, we found that within each biotype, *trans* effects were similar between conserved and not conserved TSSs (**Figure 4D**). While *trans* effect sizes were low in general, we found that conserved eRNA TSSs had the highest *trans* effect sizes (**Figure 4E**). We speculate that this may reflect the fact that enhancers are often redundant—i.e., multiple enhancers regulate the same target gene—and this may allow for them to absort *trans* effects at minimal fitness costs.

**Figure 4:**
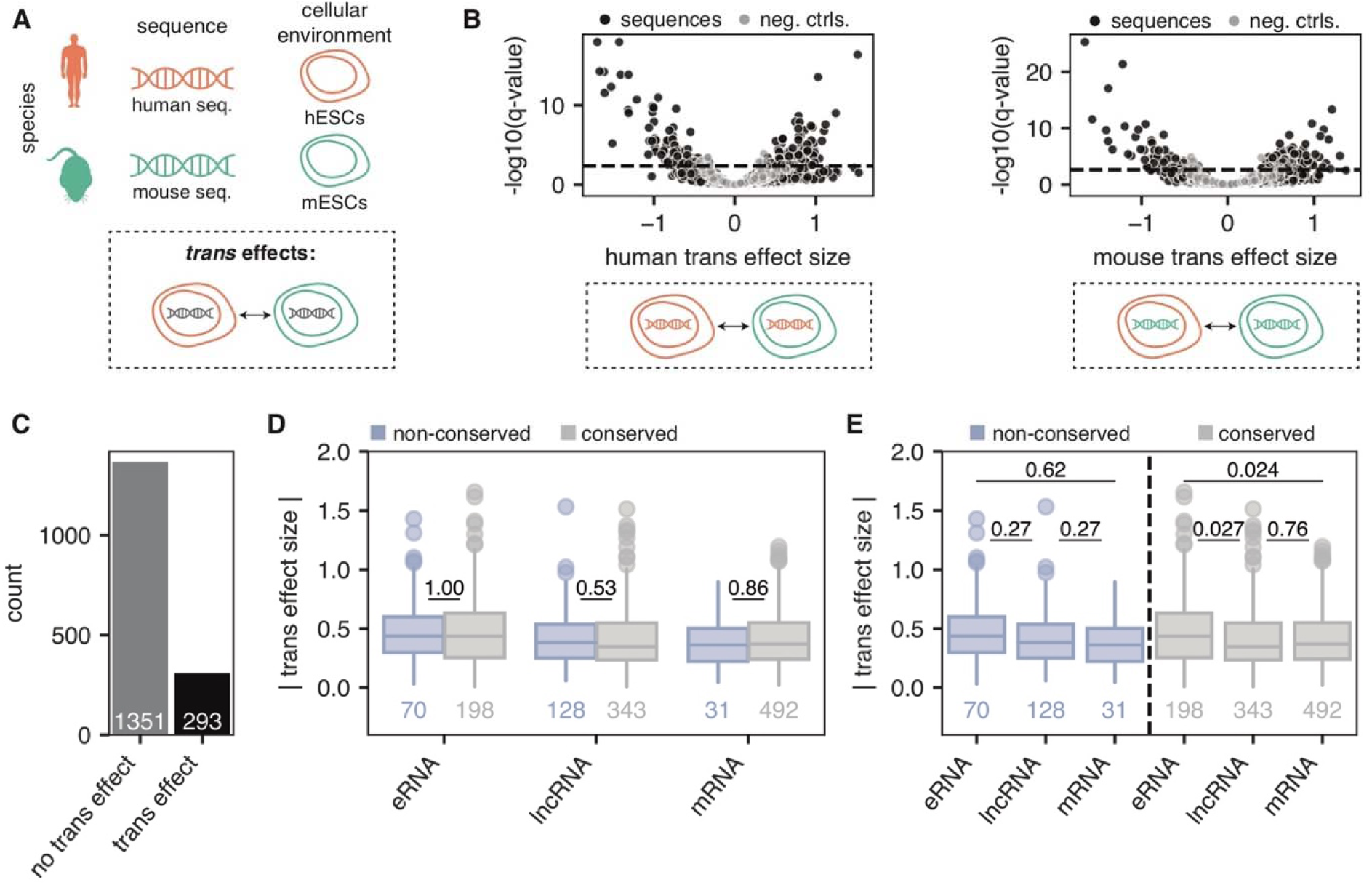
18% of orthologous regulatory elements show significant *trans* effects. **A:** Schematic depicting the definition of a *trans* effect: MPRA activity differences between hESCs and mESCs while keeping the sequence constant. **B:** Volcano plot showing the trans effect sizes (log2 foldchanges in cell type for human sequences, let and mouse sequences, right) of regulatory sequences (black) and null differential controls (gray). Horizontal line depicts an empirical FDR cut-off of 0.1, calculated using null differential controls (see methods). **C:** Count of orthologous sequence pairs with significant *trans* effects. **D:** Absolute *trans* effect sizes across biotypes, broken up into non-conserved TSSs (blue) and conserved TSSs (gray). P-values shown are from a two-sided Mann Whitney test. **E:** Same as (D), but this time comparing differences across biotypes. P-values shown are from a two-sided Mann Whitney test.

### A subset of differentially-expressed TFs are associated with *trans* effects

We next focused on identifying the TFs associated with the observed *trans* effects. We used a linear model to determine whether motif presence was significantly associated with *trans* effect sizes (see methods). After adjusting for multiple hypothesis testing, we found that 137 TFs (corresponding to 156 unique motifs) were significantly associated with *trans* effects. As motifs for different TFs can often be very similar to each other^27^ (e.g., all POU TFs share the consensus motif ATGCAAAT), we reasoned that while we found many motifs to be significantly associated with *trans* effects, only a subset of these TFs were likely driving the *trans* effect signal. To hone in on these, we performed RNA sequencing to determine differential expression of TFs between our hESCs and mESCs (see methods). We limited our analysis to the 1,032 TFs known to be one-to-one orthologs between human and mouse. Of these TFs, 661 of these TFs were expressed in either hESCs or mESCs and 428 were significantly differentially expressed (absolute log2 fold-change ≥ 1 and FDR < 0.01) between hESCs and mESCs (**Figure 5A; Supplemental Table S6**). Of the 120 TFs we found to be significantly associated with *trans* effects, 67 were one-to-one orthologs differentially expressed between hESCs and mESCs (**Supplemental Figure S11**). We reasoned that TFs likely driving *trans* effects would match in the direction of their differential expression and the direction of their *trans* effects. Of the 67 aforementioned TFs, 44 (66%) agreed in the directionality of their differential expression and *trans* effect enrichment (**Figure 5B**). These included both constitutively-active TFs (e.g. SP1, ARNT) as well as tissue-specific TFs (e.g. immune factor BACH2, developmental regulator POU2F3/OCT11) (**Figure 5C**).

**Figure 5:**
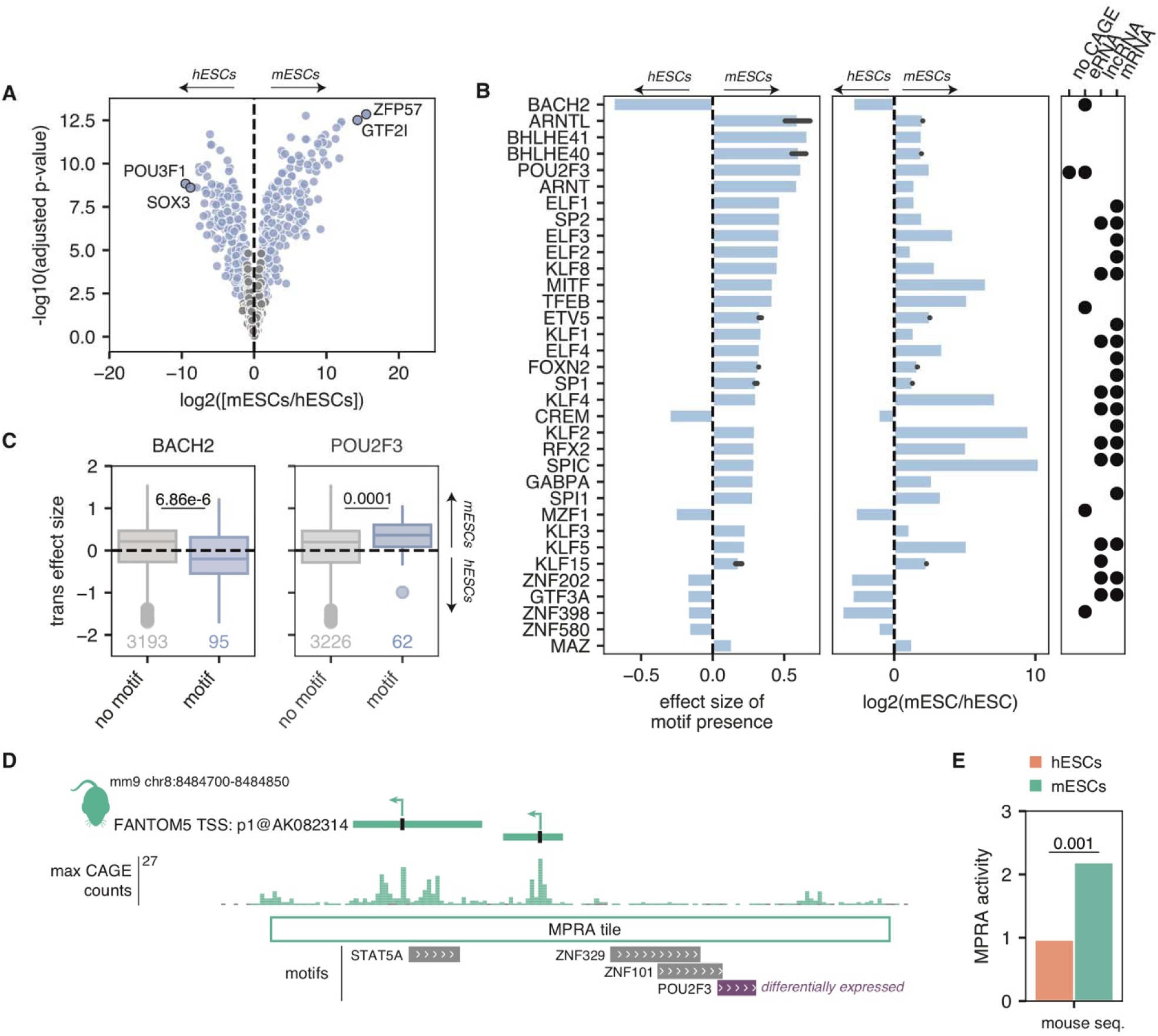
*Trans* effects are associated with a subset of differentially-expressed TFs. **A:** Volcano plot showing the differential expression of orthologous human and mouse TFs in hESCs and mESCs. Blue dots indicate significantly differentially expressed TFs (FDR < 0.01 and absolute log2 foldchange ≥ 1). The top 2 most differentially expressed TFs in either direction are highlighted. **B:** Plot showing the activating motifs significantly associated with *trans* effects (FDR < 0.05) that are also differentially expressed between hESCs and mESCs in the expected direction. Left: effect size associated with motif enrichment. Motifs that are associated with sequences more highly expressed in mESCs are > 0, and those associated with sequences more highly expressed in hESCs are < 0. Middle: log2 foldchange in expression via RNA-seq. Right: enrichment of a given TF motif across biotypes, as determined by a Hypergeometric test. Black dots denote significant enrichment (FDR < 0.05). **C:** Relationship between *trans* effect sizes and the BACH2 and POU2F3 motifs. *Trans* effect sizes for sequences with motif and without motif. *Trans* effect sizes > 0 indicate higher activity in mESCs while effect sizes < 0 indicate higher activity in mESCs. P-values shown are from two-sided Mann Whitney tests. **D:** Genome browser screenshot of an example locus showing a *trans* effect. Gray motifs correspond to TFs that are not differentially expressed between hESCs and mESCS; the purple motif, POU2F3, is differentially expressed. **E:** MPRA activities in hESCs (orange) and mESCs (green) for the mouse locus shown in D. P-value shown is the FDR value calculated by MPRAnalyze.

An example of a *trans* effect can be seen at the promoter of the uncharacterized mouse lncRNA AK082314 (**Figure 3D**). This region harbors 4 motifs for 4 TFs: STAT5A, ZNF329, ZNF101, and POU2F3. Of these 4 TFs, the only one that is differentially expressed between hESCs and mESCs is POU2F3, which is expressed ~5-fold more highly in mESCs than hESCs. Consistent with this, in our MPRA, the AK082314 promoter shows significantly higher activity in mESCs than in hESCs. Collectively, our results show that we can pinpoint a subset of TFs that may be driving *trans* effects between hESCs and mESCs.

### Co-occurrence of *cis* and *trans* effects in opposite directions are rare at individual regulatory elements

*Cis* and *trans* effects can co-occur, and previous gene-based studies have shown an excess of *cis* and *trans* effects occurring in opposite directions^6,12^. These so-called “compensatory” *cis-trans* effects help to stabilize gene expression over evolutionary time. It is unclear, however, whether the observed compensation between *cis* and *trans* effects occurs at the individual regulatory element level, or whether the compensation occurs primarily across different regulatory elements that regulate the same target gene^4^. We therefore sought to examine the extent of *cis-trans* compensation occurring within individual regulatory elements.

Of the 794 regulatory element pairs with either a *cis* or a *trans* effect, we found that 159 (20%) showed both *cis* and *trans* effects (odds = 2.01, p = 8.6 × 10^−8^, Fisher’s exact test). We then determined how often the co-occurrence of *cis* and *trans* effects was compensatory (i.e., the two effects were in opposite directions—for example, **Figure 6A** depicts a regulatory element pair with a *cis* effect showing that the mouse sequence is more active, but a *trans* effect showing that the human environment results in higher activity) or “directional” (i.e., the two effects were in the same direction—for example, **Figure 6B** depicts a regulatory element pair where the *cis* and *trans* effects are both higher for the human sequence and cell type, respectively). Surprisingly, we found that the majority (60%, p = 0.017, binomial test) of *cis-trans* co-occurrences were directional. This was driven by an excess of directional effects at non-conserved TSSs—especially eRNAs (**Figure 6C**). Conversely, conserved lncRNA and mRNA TSSs showed mostly compensatory effects. Thus, whereas regulatory element turnover between human and mouse is associated with directional *cis-trans* effects, regulatory element conservation is associated with *cis-trans* compensation.

**Figure 6:**
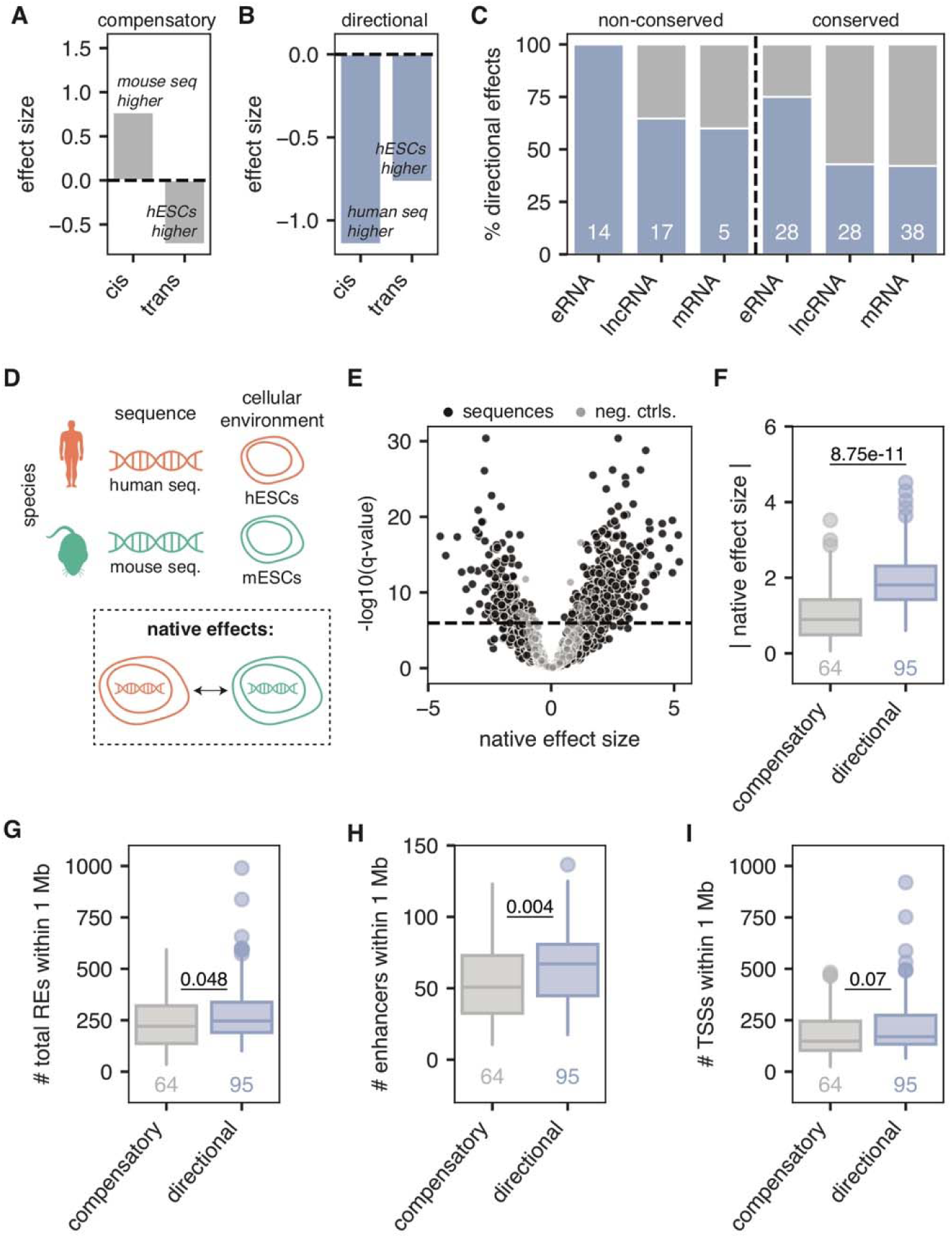
Only 40% of regulatory pairs show evidence of compensation between *cis* and *trans* effects. **A-B:** Example of a compensatory *cis-trans* effect (A) and a directional *cis-trans* effect (B). Effect sizes > 0 indicate higher activity in the mouse sequence or cellular environment whereas effect sizes < 0 indicate higher activity in the human sequence or cellular environment. **C:** Percent of regulatory element pairs across biotypes with directional *cis/trans* effects. Only pairs with both *cis* and *trans* effects are considered and the number in each group are shown. **D:** Schematic showing overview of how native effects are defined **E:** Volcano plot of native effect sizes for orthologous regulatory element pairs (black) compared to null differential controls (gray). Horizontal line depicts an empirical FDR cut-off of 0.1, calculated using null differential controls (see methods). **F:** Absolute native effect sizes for sequences showing compensatory *cis-trans* effects compared to directional *cis-trans* effects. P-value shown is from a two-sided Mann Whitney test. **G-I:** Number of additional FANTOM5 regulatory elements within a 1 Mb window of a given regulatory element, broken up by whether they show compensatory *cis-trans* effects or directional *cis-trans* effects. P-values shown are from a two-sided Mann Whitney test. G: total regulatory elements (enhancers and promoters), H: enhancers only, I: promoters only.

We next wondered whether the regulatory elements that show *cis-trans* compensation show evidence of stabilized activity between species. To this end, we examined the activity of regulatory element pairs in their native environments—human sequences in hESCs and mouse sequences in mESCs. We reasoned that if compensatory *cis-trans* effects stabilize regulatory element activity, we should see that regulatory elements in their native environments show virtually equal MPRA activities. To this end, we quantified said “native effects” between regulatory element pairs (**Figure 6D-E**). Indeed, regulatory element pairs showing compensatory *cis-trans* effects showed very low differences in native activity, whereas regulatory element pairs showing directional *cis-trans* effects showed large differences in native activity (**Figure 6F**). Thus, quantitative regulatory element activity levels are stabilized and destabilized by compensatory and directional *cis-trans* effects, respectively.

Recent work has shown that genes regulated by larger numbers of regulatory elements tend to have more stable transcription throughout evolution^13^. We therefore hypothesized that perhaps regulatory elements lacking redundancy (i.e., having fewer “partner” regulatory elements that regulate the same target gene) may show more evidence of *cis-trans* compensation than regulatory elements with high redundancy, which show more inter-element compensation. To test this, for each regulatory element, we counted the number of additional FANTOM5 regulatory elements (both TSSs and enhancers) that lied within a 1 megabase window surrounding it. We found that less redundant regulatory elements have more *cis-trans* compensation than highly redundant regulatory elements (**Figure 6G**). Moreover, this effect was driven by the number of nearby enhancers (**Figure 6H**) and not by the number of nearby promoters (**Figure 6I**). Thus, regulatory elements surrounded by few nearby enhancers tend to show compensatory *cis-trans* effects. Collectively, our results support a model whereby compensation between *cis* and *trans* effects within an individual regulatory element is more likely to occur at less redundant regulatory elements, perhaps because in these regions, there is less opportunity for inter-element compensation.

## DISCUSSION

In this work, we sought to characterize the mode underlying the evolution of individual regulatory elements that are orthologous between human and mouse by focusing on enhancers, lncRNA promoters, and mRNA promoters. Overall, we find that *trans* effects are less common and generally weaker than *cis* effects across all regulatory elements. These results are consistent with the prevailing model where *cis* effects preferentially accumulate between species, likely because *trans* effects result in more deleterious pleiotropic side-effects that are selected against^6^. We also see interesting differences between biotypes. While *cis* effect sizes are generally uniform across conserved eRNAs, lncRNAs, and mRNAs (**Figure 2E**), *trans* effects are highest in conserved eRNAs (**Figure 4E**). This suggests that the evolutionary trajectory of conserved lncRNAs is more similar to that of conserved mRNAs, whereas eRNAs behave as a separate group. Indeed, the larger accumulation of *trans* effects in eRNAs is likely due to their higher redundancy. Finally, the high resolution of our assay allowed us to identify 44 TFs that are associated with *trans* effects. Future work aimed at understanding species-level differences between human and mouse ESCs could use this set of 44 TFs as a starting point.

Previous studies have found that when *cis* and *trans* effects co-occur at the same gene, they are more often compensatory (i.e., act in different directions) than directional (i.e., act in the same direction)^6,12^, and are driven by stabilizing selection on transcript levels. However, when assessing *cis-trans* contributions at regulatory elements rather than genes, we do not find evidence of an excess of *cis-trans* compensation. Indeed, we only find evidence of enrichment for *cis-trans* compensation at conserved lncRNA and mRNA TSSs but not at eRNAs or non-conserved TSSs (**Figure 6C**). Similarly, a recent publication showed that, when looking at TF binding across different mouse strains, *cis-trans* compensation was only enriched in conserved TF binding sites^28^. Thus, while natural selection acts to stabilize the functional product of the gene—the transcript level—the evolutionary trajectories of individual regulatory elements are not as straightforward. Indeed, recent work has shown that ensembles of highly redundant enhancers are often poorly conserved, despite stable expression of their target gene throughout evolution^13^. Such data is supportive of a model wherein regulatory elements can undergo evolutionary flux and compensate for one another over time. Along these lines, here we find that highly redundant regulatory elements are less likely to show compensatory *cis-trans* effects than regulatory elements acting in smaller numbers (**Figure 6H**). Collectively, our data are consistent with the idea of inter-enhancer compensation. We propose that when regulatory elements are highly redundant, inter-enhancer compensation between *cis* and *trans* effects dominates, but when regulatory elements are less redundant, compensation between *cis* and *trans* effects can occur at the individual regulatory element level (**Figure 7**).

**Figure 7:**
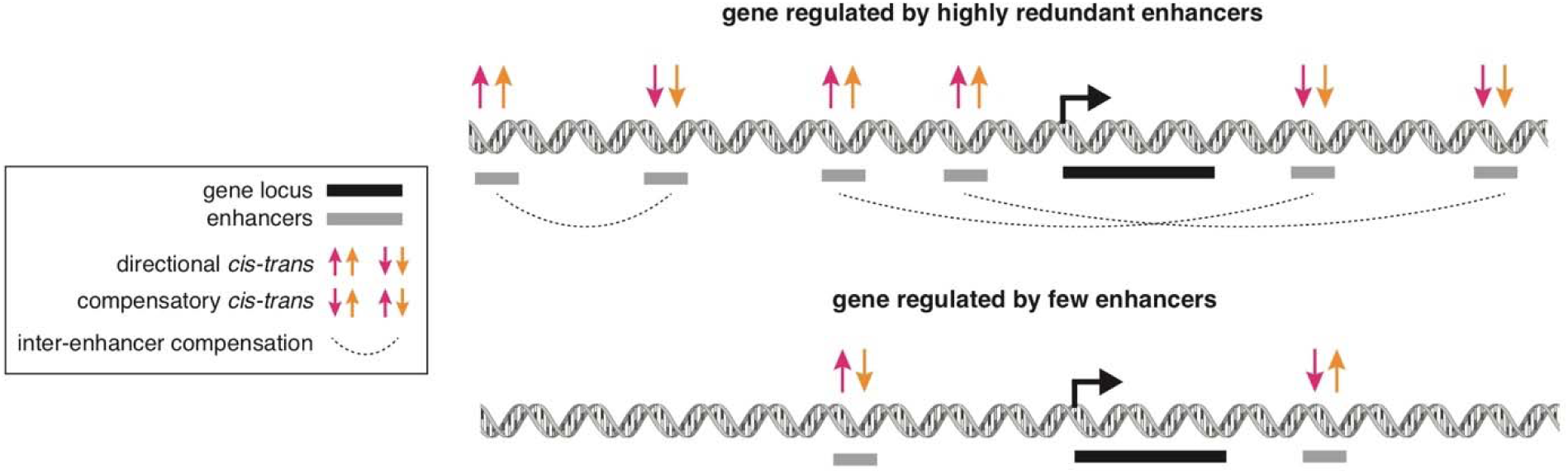
Model of inter-vs. intra-enhancer compensation. Top: at a gene regulated by many redundant enhancers, enhancers are free to show directional *cis-trans* effects because there is ample opportunity for crosstalk between enhancers, leading to inter-enhancer compensation. Bottom: at a gene regulated by very few enhancers, individual enhancers show compensatory *cis-trans* effects (intra-enhancer) because there is less opportunity for crosstalk between enhancers.

In this study we sought to perform an unbiased assessment of *cis* and *trans* effects between human and mouse across a variety of biotypes. To this end, we leveraged MPRAs to systematically test the contribution of *cis* and *trans* effects to the evolution of thousands of regulatory elements. While the use of MPRAs is extremely powerful, it also has some limitations. For example, we could only study a subset of all existing regulatory elements in the human and mouse genomes. However, the sequences that we tested were carefully selected in an unbiased manner so that they would be representative of regulatory elements genome-wide. Another limitation of our approach is that we only assessed two species in one cellular background (ESCs). Although gene expression between hESCs and mESCs is similar in general, distinct differences between the two cell lines exist^29^. Moreover, whether the known differences between hESCs and mESCs are reflective of differences in isolation and culture conditions^30^ or underlying species-specific biology remains controversial^31^. Future work is needed to assess whether similar patterns exist in other tissues, and the extent to which these patterns may affect fundamental biological processes in a species-specific manner. Nevertheless, our work has characterized the baseline to which information from other tissues and other species can be added in order to gain a more complete picture of the evolution of regulatory elements.

In summary, we find that the mode of evolution can differ at different classes of regulatory elements. Notably, we find that compensation between *cis* and *trans* effects at individual regulatory elements is rare, supporting the idea that compensation across regulatory elements—rather than within individual regulatory elements—is a widespread feature of mammalian genomes^13^. Future work should focus on integrating the mode of evolution at individual regulatory elements with target transcript expression levels across species. Collectively, our results underscore the importance of examining the role of individual regulatory elements in the evolution of gene expression.

## METHODS

### TSS selection and biotype assignment

To assign accurate TSSs to genes, we intersected human and mouse GENCODE genes^32^ (v19 in human and vM13 in mouse) with FANTOM5 TSSs^21^ in both species. Specifically, we found the closest FANTOM5 TSS (on the same strand) within +/−1000 bp of the GENCODE-annotated TSS. We classified any gene having a GENCODE gene_type of “protein_coding” as an mRNA. We classified any gene included in the GENCODE long_noncoding_RNAs gtf file as a lncRNA, provided it showed no evidence of a conserved open reading frame (PhyloCSF^33^ ORF score < 0 and branch length < 0.1). We classified any FANTOM5-annotated “robust” enhancers^34^ as eRNAs, and used both the sense and antisense TSS provided by FANTOM5. More details are available in the Supplemental Methods (TSS selection and biotype assignment section).

### Sequence orthology assignment

To determine sequence orthologs, we first mapped human TSSs (hg19) to mouse (mm9) and vice versa using the liftOver program with the parameter minMatch = 1. We then reciprocally mapped the lifted-over TSSs back to their original species, and required that they map to the exact same original TSS nucleotide. As FANTOM5 enhancers have two TSSs, we required that both TSSs reciprocally map in order to consider an enhancer a sequence ortholog.

### Conserved TSS assignment

To determine conserved vs. non-conserved TSSs, we intersected the lifted-over TSSs with the maximum CAGE read coverage in that species (ctssTotalCounts bigwig files downloaded from the FANTOM5 data hub^35^). We determined a TSS to be conserved if the region immediately surrounding the TSS (+/−50 bp) contained ≥ 10 maximum CAGE reads. As enhancers have two TSSs, if either of the TSSs intersected ≥ 10 maximum CAGE reads, we considered it conserved.

### MPRA sequence pair selection

We required all sequence pairs in the MPRA library to have a either an annotated CAGE peak in human or mouse that is expressed above background in either hESCs or mESCs (≥ 0.024 normalized counts in hESCs and ≥ 0.022 normalized counts in mESCs, **Supplemental Figure S2**). We included all lncRNAs (and their orthologous sequences) that met this threshold in the pool. We randomly selected the remaining biotypes in roughly equal numbers, given that they met this expression threshold. As eRNAs have two TSSs, we included both of its TSSs and both of its TSSs’ orthologous sequences in the pool. Exact numbers of each biotype in the MPRA can be found in **Supplemental Table S1** and the list of regulatory elements included in the MPRA can be found in **Supplemental Table S2** and. More details can be found in the Supplemental Methods (MPRA sequence pair selection section).

### MPRA oligonucleotide design

Each oligonucleotide we designed was 200 bp long, containing 144 bp of regulatory sequence, an 11 bp barcode, and 45 bp of sequence necessary for cloning. For each TSS selected above, we included two 144 bp tiles: one directly surrounding the TSS (−114/+30 bp) and one slightly upstream of the TSS (−228/-84 bp) (**Supplemental Table S3**). We then generated 1,622 random 144bp sequences to serve as negative controls. We also tiled across the CMV promoter in 144 bp segments to create 4 positive control tiles. We assigned TSS regions 13 barcodes, random sequences 3 barcodes, and CMV sequences 60 barcodes (**Supplemental Table S4**). More details can be found in the Supplemental Methods (MPRA oligonucleotide design section).

### MPRA cloning, transfection, and sequencing

Twist Bioscience synthesized the oligo pool, which we then cloned as previously described^18^ into plasmids to generate a library of constructs where the regulatory sequence is upstream of a reporter gene (here, GFP) that is upstream of a unique barcode.□ We assayed the initial representation of barcodes using high-throughput DNA-sequencing. We transfected these constructs into live cells and performed three biological replicates each in hESCs (HUES64 cells) and mESCs (derived from mouse blastocysts^36^) corresponding to three consecutive passages (**Supplemental Figure S3**). We isolated RNA and assayed barcode expression by high-throughput RNA-sequencing. More details can be found in the Supplemental Methods (MPRA cloning, transfection, and sequencing section).

### MPRA Analysis

All code to reproduce analyses is available at https://github.com/kmattioli/2019__cis_trans_MPRA.

### Quantifying MPRA activity

After trimming and quality filtering DNA and RNA reads, we mapped exact matches to known barcodes and 10 upstream constant nucleotides of GFP. We only measured sequences that had at least 50% of their barcodes represented at ≥ 10 counts in the input DNA library. We used the R package MPRAnalyze^23^ to quantify MPRA activities for each sequence in each condition using the program’s “quantification” mode. We used our randomly-generated sequences as the background null distribution, as the majority of these sequences should not induce transcription. More details can be found in the Supplemental Methods (quantifying MPRA activities section).

### Calculating differential MPRA activity

After quantifying MPRA activity and assigning 1 tile to each sequence pair (**Supplemental Figure S5**), we used MPRAnalyze^23^ to perform differential activity analyses using the program’s “comparison” mode. In comparison mode, as the null hypothesis is not the lack of transcription but the lack of differential transcription, we used down-sampled barcodes corresponding to identical CMV sequences as the background null distribution. In each of the 5 models (*cis* effects in hESCs, *cis* effects in mESCs, *trans* effects of mouse sequences, *trans* effects of human sequences, and native effects), we tested whether the full model was a better fit than an intercept-only model using a likelihood ratio test. More details can be found in the Supplemental Methods (calculating differential MPRA activity section).

### Calling significant differential effects

We considered sequences to have significant *cis, trans*, or native effects if the q-value calculated by MPRAnalyze was less than the q-value that resulted in < 10% of negative controls being called significant, which is effectively an empirical FDR of 0.1 (**Supplemental Figure S6**). We also required effect sizes to be higher than the minimum significant null differential control effect size (**Supplemental Figure S7**). We assigned each sequence pair one *cis* and *trans* effect size: we used the maximum *cis* or *trans* effect size between the two models (hESCs/mESCs for *cis* and human/mouse for *trans*) unless the effect was only significant in one model, in which case used the corresponding significant effect size. More details can be found in the Supplemental Methods (Calling significant differential effects section).

### Motif mapping

We used a curated list of human TFs defined by Lambert et al^27^. We then used the CisBP^38^ position-weight matrices designated by Lambert et al to be the “best” motifs for each of these TFs. In total, this list contained 1360 motifs corresponding to 1104 unique TFs. We mapped these motifs in both human sequences and mouse sequences using the FIMO program from the MEME suite with default parameters^39^.

### Finding motifs predictive of MPRA activity

For each motif, we fit a linear model to mean MPRA activity across all sequences as follows:

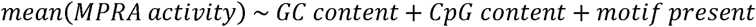

and determined whether the binary *motif present* indicator explained significantly more of the variance than a reduced model without the indicator using a likelihood ratio test (**Supplemental Table S5**). We used the Python statsmodels^40^ package to run all linear models. More details can be found in Supplemental Methods (Finding motifs predictive of MPRA activity section).

### Finding motifs associated with *cis* and *trans* effects

For each motif, we fit a linear model to absolute *cis* effect sizes across all sequence pairs as follows:

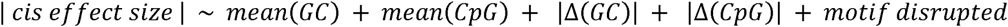

and determined whether the *motif disrupted* parameter (indicating whether a motif was present in only one of the two paired sequences) was significant (FDR < 0.05).

For each motif, we fit a linear model to *trans* effect sizes across all sequences as follows:

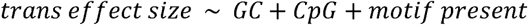

and determined whether the *motif present* parameter was significant (FDR < 0.05). More details available in Supplemental Methods (Finding motifs associated with *cis* and *trans* effects sections).

### RNA-seq of hESCs and mESCs

We sequenced both untransfected and transfected hESCs and mESCs. We extracted RNA from TRIzol using standard protocols and used the Illumina TruSeq kit (non-stranded) to create polyA+ libraries from total RNA. We measured library concentration using the Qubit dsDNA HS Assay kit (Thermo Fisher Scientific), and ran all of the libraries on a Bioanalyzer (Agilent) to assess purity and fragment size, and sequenced on a HiSeq 2500 at Harvard University’s Bauer Sequencing Core (75 bp paired end).

### RNA-seq analysis

We aligned reads to either hg19 or mm10 using Hisat2^41^. We used FeatureCounts to count reads aligning to genes in either GENCODE v25 (human) or GENCODE vM13 (mouse)^42^. We downloaded orthologous genes between human and mouse from Ensembl (version 96)^43^, and removed any orthologs classified as “many-to-many”. We quantified gene expression in each transfected sample using DESeq2^44^. To find differentially expressed genes, we used the edgeR-limma pipeline^45^ (filtering out any genes with cpm < 1) to model paired samples (transfected and untransfected) and control for transfection status.

## ACKNOWLEDGEMENTS

We thank Veronika Akopian for help with HUES64 cell culture, Abigail Groff for providing mESCs, Jordan Lewandowski for help with mESC cell culture, and Martha Bulyk for helpful discussions. M.M. was a Gilead Fellow of the Life Sciences Research Foundation during part of the project. K.M. was a National Science Foundation Graduate Research Fellow under grant no. DGE1144152 during the majority of the project. J.L.R. is an HHMI faculty scholar.

## AUTHOR CONTRIBUTIONS

K.M. and M.M. designed the project and wrote the manuscript. K.M. designed the oligonucleotide libraries and performed all MPRA computational analyses. W.O. analyzed FANTOM5 CAGE data and performed RNA-seq analysis. C.G. and P.G.M. performed the MPRA experiments. C.G. performed RNA-seq. D.A. assisted with cell culture. J.L.R. contributed to the project design and discussion. All authors have read and approved the manuscript for publication.

## DATA ACCESS

The MPRA sequencing data and genomic RNA-seq data from this study have been submitted to the NCBI Gene Expression Omnibus (GEO; http://www.ncbi.nlm.nih.gov/geo/) under accession number GSE140574. All scripts required to reproduce this work are available on GitHub at https://github.com/kmattioli/2019__cis_trans_MPRA

