## Supplemental Materials for "Analysis of regulatory element evolution between human and mouse reveals a lack of *cis-trans* compensation"

### **Supplemental Methods**

#### ***TSS selection and biotype assignment***

**TSS selection.** The 5’ ends of genes are often misannotated, and these annotations can be improved by CAGE-seq, which sequences the 5’ ends of RNA more accurately than traditional RNA-seq^1^. The FANTOM5 consortium has performed CAGE-seq across thousands of human and mouse samples, and identified accurate 5’ ends of protein-coding genes and non-coding RNAs at single-nucleotide resolution. Thus, to get an accurate list of TSSs for genes, we assigned FANTOM5 TSSs to genes in both human and mouse. We intersected FANTOM5 TSSs defined as “robust” with the GENCODE-annotated gene TSSs (comprehensive annotation, v19 in hg19 for human and vM13 in mm10 for mouse). In order to intersect the mouse genes (mm10) with FANTOM5 CAGE data (mm9), the vM13 GENCODE transcripts were aligned to the mm9 genome using Crossmap^2^. We assigned the closest FANTOM5 TSS on the same strand within +/- 1000bp of the GENCODE-annotated gene start to GENCODE genes. To select transcribed enhancer TSSs, we selected the “robust” enhancers identified by the FANTOM5 consortium in both species^3^. These TSSs are defined by CAGE-seq and a sense and antisense TSS are annotated for each enhancer, both of which we selected.

**Biotype assignment.** For both species, we assigned any genes that had a gene_type of “protein_coding” in the comprehensive GENCODE annotation gtf file (v19 for human and vM13 for mouse)^4^ to be mRNAs. We assigned any genes included in the long_noncoding_RNAs.gtf file provided by GENCODE to be lncRNAs, but removed those had conserved open reading frames (ORFs). We found conserved ORFs using the PhyloCSF program^5^, using an alignment of 29 mammals and the parameters --frames=3, --orf=ATGStop, and --minCodons=3. We re-classified any lncRNA with a maximum ORF score of > 0 and a maximum branch length of > 0.1 to be an mRNA if the ORF was longer than 150 amino acids or “other” if the ORF was shorter than 150 amino acids. As mentioned above, we assigned any enhancers annotated in the “robust” FANTOM5 annotation files in both species to be eRNAs.

#### ***MPRA library design***

**MPRA sequence pair selection.** To select sequence pairs to add to the MPRA, we focused only on sequence orthologs. We then determined expression of TSSs in hESCs and mESCs based on the FANTOM5 normalized expression values for hESCs (*sample IDs: CNhs14067, CNhs14068, CNhs13964, CNhs11917, CNhs12824, CNhs12837, CNhs13694, CNhs13695, CNhs13738*) and mESCs (*sample IDs: CNhs14104, CNhs14109, CNhs14098, CNhs14099, CNhs14100, CNhs14091, CNhs14092, CNhs14093*). We then subset sequences to those expressed above background in either hESCs or mESCs (≥ 0.024 normalized counts in hESCs and ≥ 0.022 normalized counts in mESCs, **Supplemental Figure S2**). We included all human or mouse intergenic lncRNAs (≥ 1000 bp away from any annotated protein-coding genes) that met this threshold. We then randomly sampled all other lncRNAs as well as mRNAs in roughly equal numbers. We randomly sampled roughly half as many eRNAs, as each eRNA has two TSSs. For each sequence sampled, we included its orthologous region in the pool. After designing the pool, we re-classified some lncRNAs as micropeptides given that they contained conserved ORFs (see above). We assigned these as a biotype of “other”. The final counts of biotype pairs in the MPRA library can be found in **Supplemental Table S1** and the full list of sequences included in the MPRA library can be found in **Supplemental Table S2**.

**MPRA oligonucleotide design.** We designed one 240,000 oligonucleotide (oligo) pool of 200 bp and ordered the pool from Twist Bioscience. We assigned TSSs each 13 barcodes, random negative control sequences 3 barcodes, and positive control sequences (tiled regions of the CMV promoter) 60 barcodes. The final oligo distribution can be found in **Supplemental Table S4**. All oligos contained universal primers, two restriction enzyme sites, an 11 bp barcode, and the regulatory sequences of interest as follows^6^:

**5’**---**universal primer 1** (16 bp)---**variable sequence** (144 bp)---**XbaI** (6bp)---**KpnI** (6bp)---**barcode** (11 bp)---**universal primer 2** (17 bp)---**3’**

**Universal primer 1:** ACTGGCCGCTTCACTG

**Universal primer 2:** AGATCGGAAGAGCGTCG

#### ***MPRA cloning, transfection, and sequencing***

**ePCR amplification of oligo pools.** The synthesized oligo pool was amplified by emulsion-PCR (ePCR, Micellula DNA Emulsion & Purification Kit, Chimerx), according to the manufacturers’ instructions. ePCR primers harboring Sfi I restriction sites were designed to add for subsequent cloning purposes (5’ primer: GCTAAGGGCCTAACTGGCCGCTTCACTG; 3’ primer: GTTTAAGGCCTCCGAGGCCGACGCTCTTC). To determine the oligos representation of the ePCR-amplified oligopool (based on the unique 3’ barcode of each oligo), 1 ng of the amplified oligo pool was used as input for library preparation (see below) and sequenced on a MiSeq (SR, Illumina).

**Cloning.** The ePCR-amplified oligopools were digested with SfiI, and inserted in the multiple cloning region of an empty vector. In the first cloning step, the ligation reaction (100 ng backbone + 4x molar excess of oligopool) was transformed into 20 x DH5α tubes (ThermoScientific), spread out on 50 ampicillin LB plates, and incubated overnight at 37°C. After scraping all bacteria of each LB plate in 5 ml LB and pooling all colonies, plasmids were purified with the endotoxin-free Qiagen Plasmid Plus Maxi kit (Qiagen). The oligo representation was determined by MiSeq-sequencing. Next, the cloned oligopool was sequentially cut with Kpn I and Xba I and ligated as described above with and without a minimal promoter (5’-AGAGGGTATATAATGGAAGCTCGACTTCCAG-3’) and an ORF for GFP. Illumina sequencing determined barcode representation of the oligopools. Finally, to remove plasmids without inserted oligos, the oligopools were digested with Kpn I and the plasmid fractions containing oligos were size-selected by agarose gel-electrophoresis, re-ligated and cloned as described above.

**Library preparation.** Cloned oligopools (50 ng) were amplified with PfU HS DNA polymerase (Agilent) and 5 µl of 2 µM index primer (cloning step 1, universal 3’ primer: AATGATACGGCGACCACCGAGATCTACACTCTTTCCCTACACGACGCTCTTCCGATCT; Index 1 5’ primer: caagcagaagacggcatacgagatCGTGATgtgactggagttcagacgtgtgctcttccgatctACTGGCCGCTTCACTG; cloning step 2 & 3 and cDNA libraries, universal 3’ primer: AATGATACGGCGACCACCGAGATCTACACTCTTTCCCTACACGACGCTCTTCCGATCT; Index 1 5’ primer: caagcagaagacggcatacgagatCGTGATgtgactggagttcagacgtgtgctcttccgatctCGCCGCGTGGAGGAGGA, underlined nucleotides = index; PCR setting: 95°C 2 min, 95°C 30 sec, 55°C 30 sec, 72°C 15 sec [x 18-24 cycles], 72°C 10 min, 4°C hold). After 18 cycles of amplification, molarity was checked on a BioAnalyzer. Insufficient amplifications were run for 2-6 additional cycles (total of 24 cycles). Amplified libraries were then cleaned three times with AMPure beads with the following ratios according to manufacturer’s instruction: 0.6x, 1.6x, 1.0x.

**Transient transfections.** HUES64 cells were cultured in feeder-free mTeSR1 media (Stem Cell Technologies 85850) which was prepared as recommended by the manufacturer. mESC cells were derived from mouse blastocysts^7^ and were cultured in feeder-free 2i media which was prepared as previously described^7^. Tissue culture conditions were standard (5 % CO_2,_ 37°C) for both cell lines. HUES64 cells were passaged every 5-7 days whereas mESC cells were passaged every 3-5 days. For the HUES64 replicates, as transfections were done on adherent cells, which limited the number of cells per transfection, 3 independent transfections (technical replicates) were performed for each of 3 biological replicates (3 different passages). For each transfection of HUES64 cells, 2.5 million cells were transfected with 25 µg of plasmid using 60 µL of Lipofectamine Stem (Thermo Fisher Scientific). A total of 7.5 million cells were transfected for each biological replicate of HUES64. For the mESC replicates, as the transfections could be done in suspension with higher numbers of cells, single transfections of 3 biological replicates (3 different passages) were performed. For each biological replicate of mESCs, 40 µg of plasmid were transfected into 10 million cells using 100 µL of Lipofectamine 2000 (Thermo Fisher Scientific). In both cases, RNA was harvested 24-hours post-transfection and transfection efficiency was microscopically determined by GFP expression (**Supplemental Figure S3**).

**RNA extraction and cDNA library preparation.** RNA from transiently transfected hESCs and mESCs was precipitated by phenol-chloroform extraction according to standard protocols. DNase treatment (Worthington) was followed by cDNA synthesis with SuperScript III First-Strand Synthesis System (Invitrogen). cDNA was subject to library amplification as listed above (primer: cloning step 2 & 3 and cDNA libraries), and libraries were cleaned up with AMPure beads (0.6x, 1.6x, 1.0x).

**Sequencing & quality control.** Genomic DNA (input library) or polyA+ RNA was sequenced using the Illumina single end 50 bp kit. The program cutadapt ^8^ was used to remove adapters and trim bases with a Phred score lower than 20. Each filtered read was required to exactly match one of our pre-designed 11-nucleotide barcodes as well as the neighboring upstream 10 constant nucleotides of GFP (TCTAGAATTA), otherwise it was discarded.

#### ***MPRA analysis***

All scripts used to do the MPRA analysis are available at <https://github.com/kmattioli/2019__cis_trans_MPRA>. Most code is in Python, with the exception of the MPRAnalyze and DESeq2/edgeR code, which is in R.

**Data pre-processing.** We required elements to have at least 50% of their barcodes with counts ≥ 10 in the input DNA library.

**Quantifying MPRA activities.** We used “quantification mode” in the MPRAnalyze program^9^ to quantify MPRA activities for each sequence in each condition (hESCs and mESCs). We used our randomly-generated negative control sequences to serve as the background null distribution, as the vast majority of these sequences should not induce transcription. We also included positive control barcodes that were down-sampled from our 4 unique positive control sequences. As each of the 4 positive control sequences had 60 barcodes, we randomly sampled 13 of these barcodes 100 times for each sequence to create a set of 400 positive control barcode counts. We determined sequences to have significant activity if their MPRAnalyze q-value was < 0.05.

**Calculating differential MPRA activity.** We used “comparison mode” in the MPRAnalyze program to quantify differential activity. Since the null hypothesis in comparison mode is not the lack of transcription but the lack of differential transcription, we paired the down-sampled CMV barcodes used above, and used these as null differential controls. Specifically, we randomly paired sets of 13 down-sampled barcodes corresponding to the same CMV sequence 100 times for each of 4 CMV sequences to create a set of 400 null differential control sequences.

In the *cis* effects analysis, we separately tested for differential activity between human and mouse sequences in hESCs and human and mouse sequences in mESCs. In the *trans* effects analysis, we separately tested for differential activity between hESCs and mESCs for human sequences and for mouse sequences. In the native effects analysis, we tested for differential activity between human sequences in hESCs and orthologous mouse sequences in mESCs. In each of the aforementioned cases, we tested whether the defined model was a better fit than an intercept-only model using a likelihood ratio test.

**Calling significant differential effects.** We noticed that each of the 5 models (*cis* effects in hESCs, *cis* effects in mESCs, *trans* effects of human sequences, *trans* effects of mouse sequences, and native effects) found a very different portion of null differential controls as significant at an MPRA q-value alpha of < 0.05 (**Supplemental Figure S6**), indicating that each of the 5 models had varying levels of statistical power. To control for this, we found the 10^th^ percentile MPRAnalyze q-value in each model across the set of 400 null differential controls; this is an FDR of < 0.1 across all models, as it results in < 10% of null differential controls being called significant in each model.

We also noticed that a small subset of sequences with FDR < 0.1 had very low effect sizes, and so we did not consider these significant. To this end, we found the minimum effect size corresponding to significant null differential controls (FDR < 0.1 threshold) in each model. If any sequence pair had an effect size less than this effect size threshold, we did not consider it significant (**Supplemental Figure S7**).

We considered sequences to have significant differential activity if their q-values were below these model-specific FDR < 0.1 q-value thresholds and their effect sizes were above these model-specific effect size thresholds. If a sequence pair showed a *cis* effect in either model (hESCs or mESCs), we considered the pair to have a significant *cis* effect. Similarly, if a sequence showed a *trans* effect in either model (human sequences or mouse sequences), we considered the pair to have a significant *trans* effect.

As we ran two models each for *cis* and *trans* effects (*cis* in hESCs/mESCs and *trans* for human sequences/mouse sequences), we had two *cis* and *trans* effect sizes for each sequence pair. We assigned one representative *cis* effect size as follows: (1) if there was a significant *cis* effect in both hESCs and mESCs, or if there was no significant *cis* effect in both hESCs and mESCs, assign the maximum *cis* effect size of the two, (2) if there was only a significant *cis* effect in hESCs, assign the *cis* effect size calculated in hESCs, and (3) if there was only a significant *cis* effect in mESCs, assign the *cis* effect size calculated in mESCs. Similarly, we assigned one representative *trans* effect size as follows: (1) if there was a significant *trans* effect in both human and mouse sequences, or if there was no significant *trans* effect in both human and mouse sequences, assign the maximum *trans* effect size of the two, (2) if there was only a significant *trans* effect in the human sequence, assign the *trans* effect size calculated for the human sequence, and (3) if there was only a significant *trans* effect in the mouse sequence, assign the *trans* effect size calculated for the mouse sequence.

#### ***Motif analysis***

**Finding motifs predictive of MPRA activity.** We first calculated the GC content (%Gs and Cs) and CpG content (% CG dinucleotides) in each sequence. We then Box-Cox transformed the mean MPRA activity values to “normalize” the distribution, and standardized the dataset to have a mean of 0 and a standard deviation of 1. Once the data was pre-processed, then for each motif, we fit a linear model to mean MPRA activity across all sequences as follows:

$$mean\left( MPRA activity \right) \sim GC content+CpG content+motif present$$

and determined whether the binary $motif present$ indicator explained significantly more of the variance than a reduced model without the indicator using a likelihood ratio test. The reduced model of just GC and CpG content was able to explain 15.1% of variance in mean MPRA activity. We considered a motif to explain a significant amount of variance in MPRA activity if its Benjamini-Hochberg adjusted p-value was < 0.05. We also used this model to predict activating and repressive motifs. If the beta value returned for the $motif present$ indicator was positive, we considered the motif activating; if it was negative, we considered it repressive. The full list of motif results from this model can be found in **Supplemental Table S5**.

**Finding motis associated with *cis* effects.** As *cis* effects are between two sequences, we first calculated the mean GC and CpG content across each pair of sequences. We also calculated the absolute difference in GC and CpG content between each pair of sequences. We then Box-Cox transformed the absolute *cis* effect sizes to “normalize” the distribution, and standardized the dataset to have a mean of 0 and a standard deviation of 1. We then limited our model to only sequences that had significant activity in either hESCs or mESCs. To find motifs enriched in *cis* effects, for each motif, we broke sequence pairs up into three “motif classes”: those where the motif is not present in either species (“not present”), those where the motif is maintained in both species (“maintained”), and those where the motif is only present in one species (“disrupted”). We fit the following linear model:

$$\left| cis effect size \right| \sim mean\left( GC \right)+mean\left( CpG \right)+ \left| \Delta\left( GC \right) \right|+ \left| \Delta\left( CpG \right) \right|+motif class$$

and found the p-value and beta assigned to the comparison of the “disrupted” factor to the “maintained” factor within the $motif class$ parameter. We considered a motif to be significantly associated with *cis* effects if its Benjamini-Hochberg adjusted p-value was < 0.05 and it had ≥ 10 sequences with a “maintained” motif. We also required that the beta be > 0, as a negative beta would indicate that maintaining the motif was positively correlated with *cis* effects. However, all 17 significant motifs that we found indeed had beta values > 0.

**Finding motifs associated with *trans* effects.** As we did not use absolute effect sizes for the *trans* model, we did not need to Box-Cox transform the effect sizes. We standardized the dataset to have a mean of 0 and a standard deviation of 1. We then limited our model to only sequences that had significant activity in either hESCs or mESCs. To find motifs enriched in *trans* effects, for each motif, we fit the following model:

$$trans effect size \sim GC+CpG+motif present$$

and found the p-value and beta for the $motif present$ parameter. We considered a motif to be significantly associated with *trans* effects if its Benjamini-Hochberg adjusted p-value was < 0.05 and there were ≥ 10 sequences that contained the motif. As we used raw *trans* effect sizes, we could use the beta values to determine whether the motis were associated with *trans* effects that were higher in mESCs (beta > 0) or higher in hESCs (beta < 0).

### **Supplemental Tables**

**Supplemental Table S1: Count of unique orthologous regulatory element pairs included in the MPRA.** Rows denote human biotypes whereas columns denote mouse biotypes. Gray shaded squares correspond to sequences potentially analyzed in “conserved TSS” analyses whereas blue shaded squares correspond to sequences potentially analyzed in “non-conserved TSS” analyses. *eRNAs: as eRNAs have two TSSs, sometimes one TSS can map to one biotype while the second TSS maps to a different biotype; the number of unique human eRNAs is 309 and the number of unique mouse eRNAs is 377. ^†^other: biotypes not explicitly analyzed in this paper, which include micropeptide promoters (lncRNAs found to contain conserved ORFs) and multi-mapped promoters (promoters which could not be unequivocally assigned a biotype). ^††^3,327: because of the aforementioned enhancer issue, this table includes some duplicate orthologous regulatory elements. The total number of unique orthologous regulatory elements included in the MPRA is **3,275**.

|  |  | ***mouse*** | | | | | **TOTAL** |
| --- | --- | --- | --- | --- | --- | --- | --- |
|  |  | **eRNAs*** | **lncRNAs** | **mRNAs** | **other^†^** | **no CAGE activity** |  |
| ***human*** | **eRNAs*** | 255 | 3 | 6 | 5 | 97 | **366** |
|  | **lncRNAs** | 84 | 657 | 63 | 47 | 250 | **1,101** |
|  | **mRNAs** | 2 | 0 | 1,000 | 30 | 49 | **1,081** |
|  | **other^†^** | 23 | 25 | 83 | 281 | 62 | **474** |
|  | **no CAGE activity** | 89 | 134 | 43 | 39 | 0 | **305** |
|  | **TOTAL** | **453** | **819** | **1,195** | **402** | **458** | **3,327^††^** |

**Supplemental Table S2: List of regulatory elements included in the MPRA.** Table is a txt file. Detailed column descriptions provided in file.

**Supplementary Table S3: Count of elements in the oligonucleotide library, separated by species, biotype, and tile number** (tile 1: TSS-overlapping tile, tile 2: upstream tile). Tile numbers are not exactly equal because when an oligonucleotide was found to contain a restriction enzyme site, we dropped it from the library. *eRNAs: note that these numbers include both TSSs for eRNAs.

|  | ***human*** | | | ***mouse*** | | | ***TOTAL*** |
| --- | --- | --- | --- | --- | --- | --- | --- |
|  | **tile 1** | **tile 2** | **subtotal** | **tile 1** | **tile 2** | **subtotal** |  |
| eRNAs* | 641 | 628 | 1,269 | 672 | 661 | 1,333 | 2,602 |
| lncRNAs | 1,040 | 1,036 | 2,076 | 718 | 726 | 1,444 | 3,520 |
| mRNAs | 1,025 | 1,022 | 2,047 | 1,072 | 1,059 | 2,131 | 4,178 |
| other^†^ | 430 | 439 | 869 | 344 | 349 | 693 | 1,562 |
| no CAGE activity | 347 | 348 | 695 | 485 | 491 | 976 | 1,671 |
| ***TOTAL*** | | | | | | | 13,533 |

**Supplementary Table S4: Count of elements and barcodes in the oligonucleotide library.** *Unanalyzed sequences: sequences that we included in this library for a separate project that were not analyzed in this paper.

|  | **# elements** | **# barcodes** | **total oligos** |
| --- | --- | --- | --- |
| TSSs | 13,533 | 13 | 175,929 |
| Positive controls (CMV) | 4 | 60 | 240 |
| Negative controls (random sequences) | 1,622 | 3 | 4,896 |
| Unanalyzed sequences* | --- | --- | 58,588 |
| ***TOTAL*** | | | **239,653** |

**Supplemental Table S5: TF motifs and their effects on MPRA activity.** Table is a txt file. Detailed column descriptions provided in file.

**Supplemental Table S6: Orthologous TF expression in hESCs and mESCs.** Table is a txt file. Detailed column descriptions provided in file.

### **Supplemental Figures**

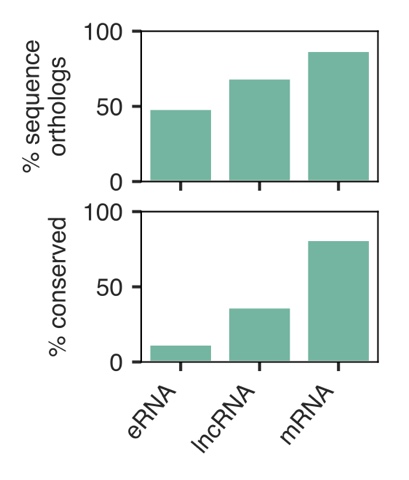

**Supplemental Figure S1: Percentage of mouse-to-human sequence orthologs and conserved TSSs broken up by biotype.** TSSs are considered conserved if the orthologous region in the other species was within 50 bp of ≥ 10 maximum CAGE reads.

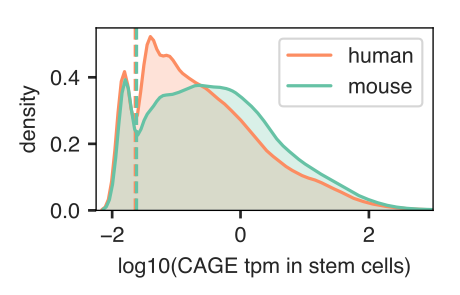

**Supplemental Figure S2: CAGE expression cut-off used to include sequences in the MPRA.** We required either the human or the mouse TSS region to overlap an annotated CAGE peak that was expressed above background in either hESCs or mESCs. We considered this background threshold (vertical dashed lines) to be at the “shoulder” of the distribution of expression values across TSSs and enhancers. In hESCs, this level was ≥ 0.024 normalized counts, whereas in mESCs, this level was ≥ 0.022 normalized counts.

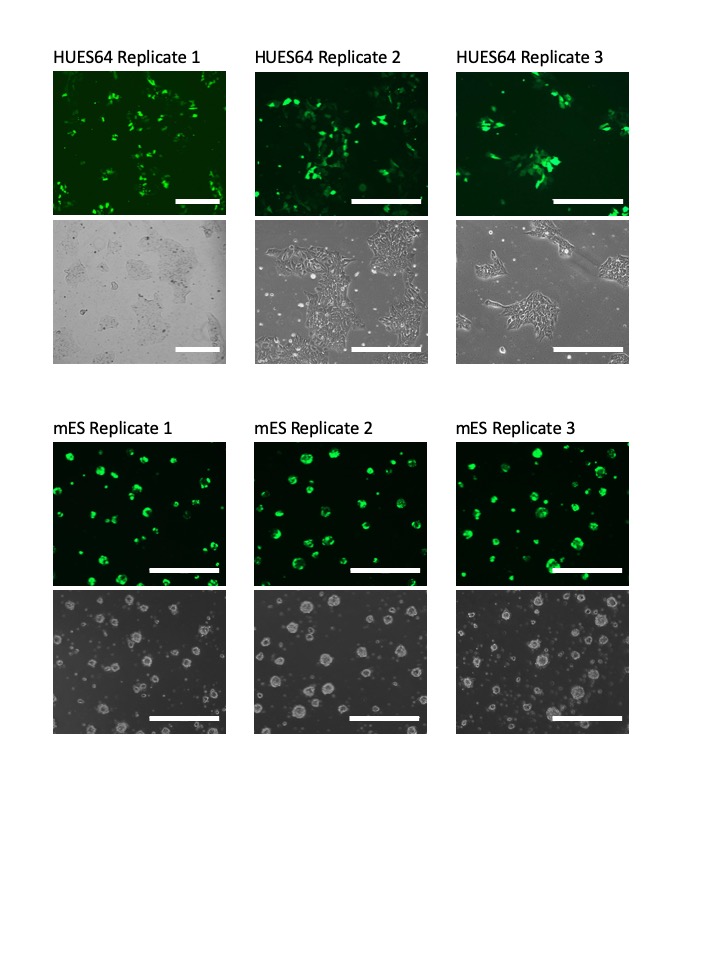

**Supplemental Figure S3: Transfection efficiencies across biological replicates.** HUES64 cells and mESCs transfected with the pMAX-GFP control plasmid (same condition as for the MPRA plasmid transfections; for HUES64: 10ug plasmid/1M cells, lipofectamineStem/DNA ratio 1:2.5, for mESCs: 4ug plasmid/1M cells, Lipofectamine 2000/DNA ratio 1:2.5). Scale bar = 400 µm for all images. (Note: replicate 1 was imaged at lower magnification.)

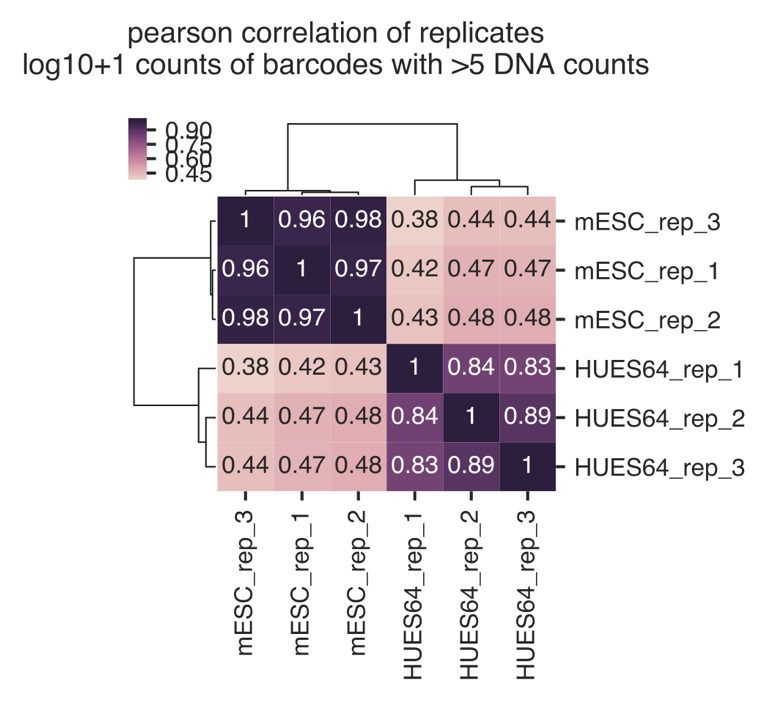

**Supplemental Figure S4: Heatmap showing correlation of barcode counts across biological replicates.** Only barcodes with at least 5 DNA counts are shown, and samples are clustered hierarchically. As expected, hESC (HUES64) replicates cluster together, as do mESC replicates, and the two species cluster separately.

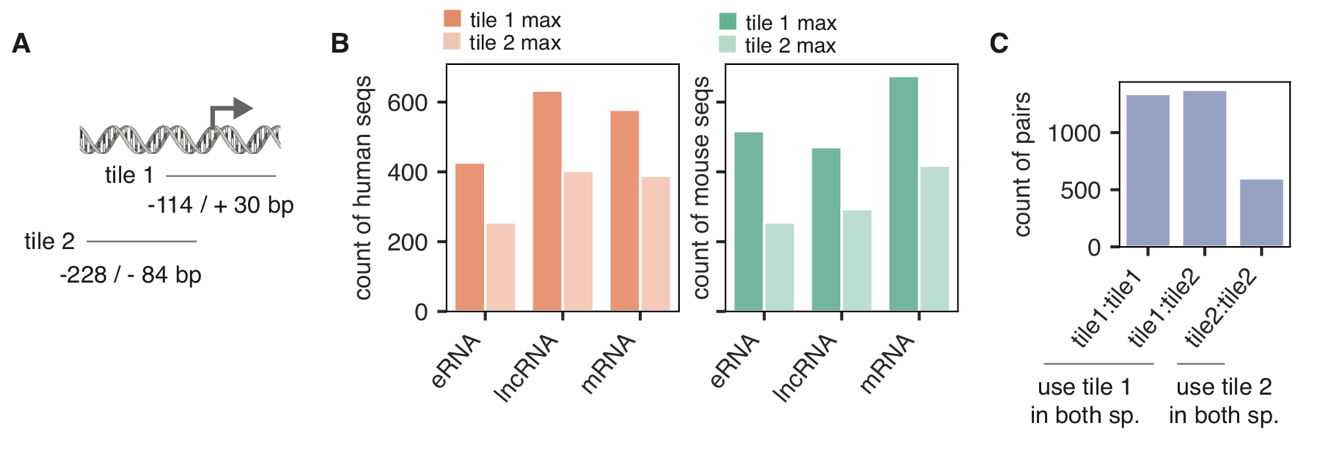

**Supplemental Figure S5: Analysis of tiles with maximum activity. A:** Schematic showing the locations of tile 1 (114 bp upstream to 30 bp downstream of the FANTOM5-defined TSS or lifted-over TSS) and tile 2 (228 bp upstream to 84 bp upstream of the FANTOM5-defined TSS or lifted-over TSS). **B:** Count of either human (left) or mouse (right) sequences showing maximum activity in either tile 1 or tile 2 across biotypes. **C:** Count of maximum tile pair designations: tile1:tile1 = tile 1 is the maximum tile in both species, tile1:tile2 = tile 1 is the maximum tile in 1 species but tile 2 is the maximum tile in the other species, tile2:tile2 = tile 2 is the maximum tile in both species. We used tile 1 in both species for tile1:tile1 and tile1:tile2, but tile 2 in both species for tile2:tile2.

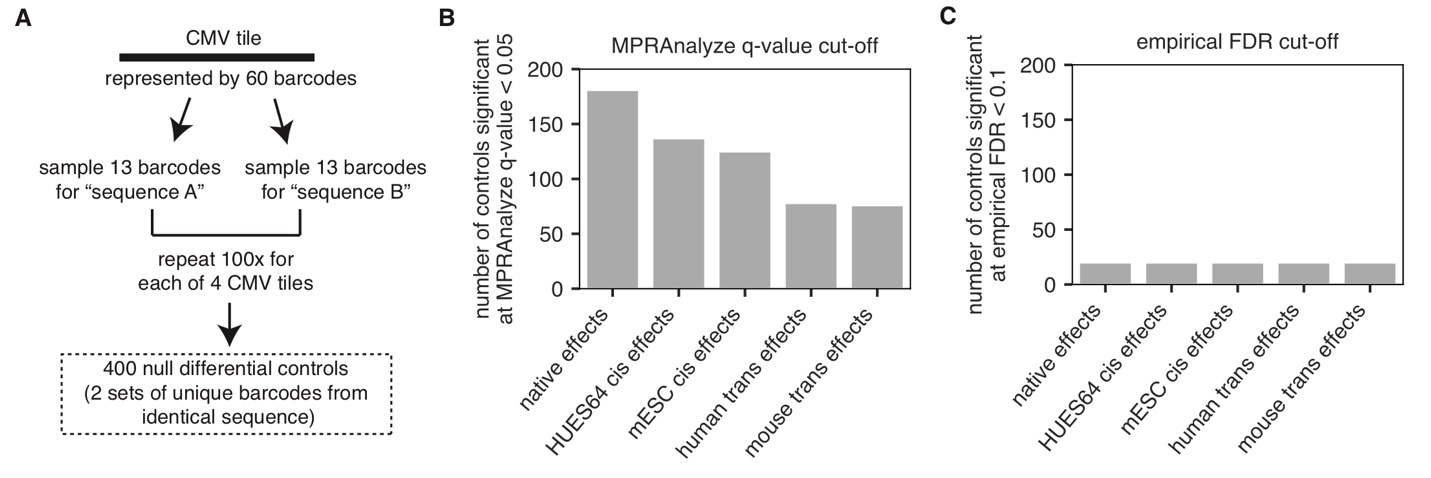

**Supplemental Figure S6: Using null differential controls to set an empirical FDR for each MPRAnalyze model. A:** Schematic showing how we created null differential controls. From each CMV tile, we sampled (without replacement) 13 barcodes to serve as “sequence A” and 13 barcodes to serve as “sequence B”, and repeated this 100 times for each of 4 CMV tiles in the library for a total of 400 null differential controls. **B:** Different MPRAnalyze models had different levels of statistical power, as observed by the number of null differential controls with MPRAnalyze q-values of < 0.05 in each model. **C:** To ensure that we had similar levels of statistical power across models, we set a model-specific q-value threshold at which point < 10% (< 40/400) null differential controls would be called as “significant” in the model.

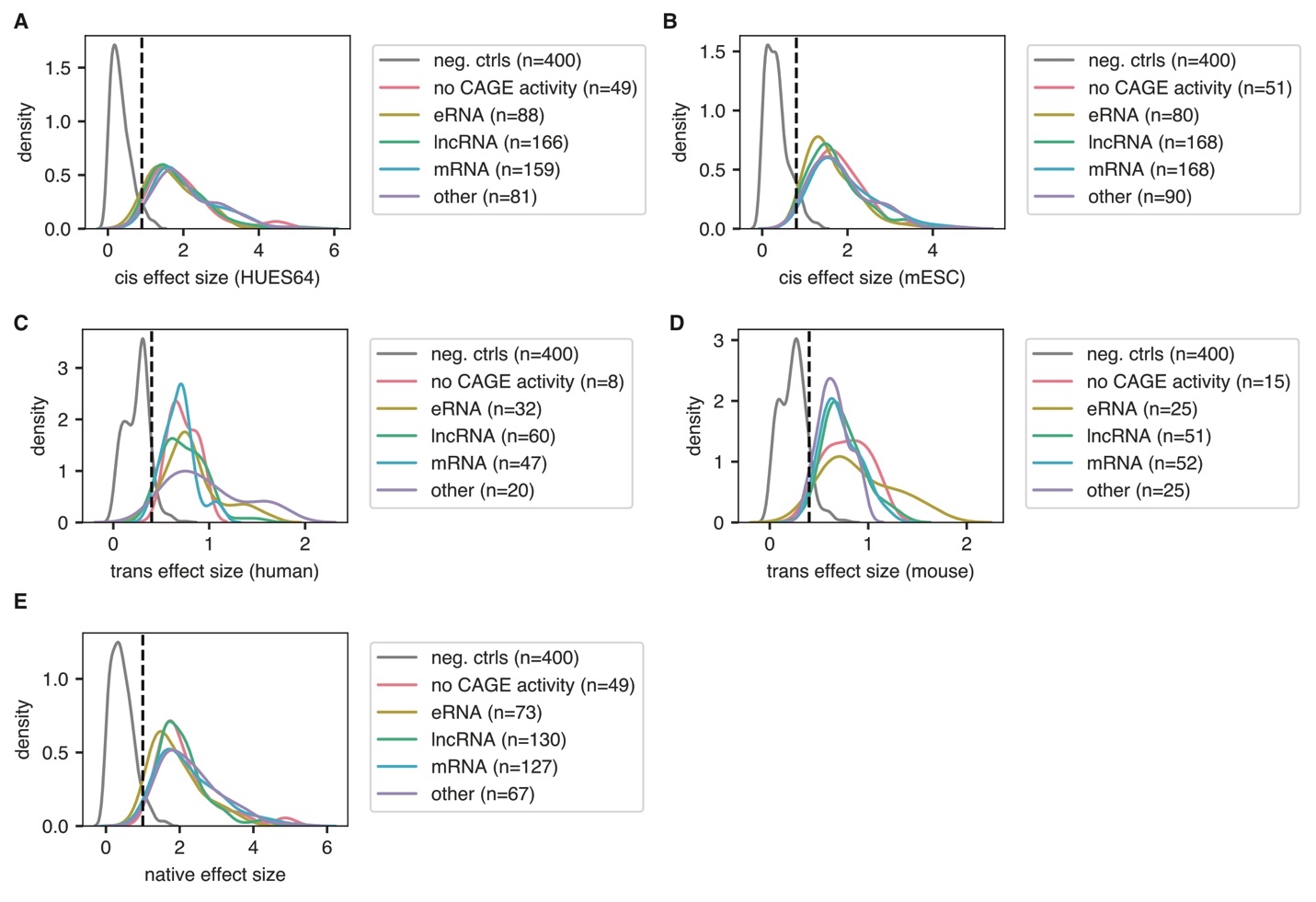

**Supplemental Figure S7: Model effect sizes for null differential controls and TSSs.** Density plot showing the absolute effect sizes calculated by MPRAnalyze in each model for sequences with significant differential activity (FDR < 0.1). Biotypes, including null differential controls (labeled “neg. ctrls”) are labeled, with the number of sequences with FDR < 0.1. As some sequences with significant differential activities had very low effect sizes, we set a model-specific effect size cut-off. We required significant sequences to have absolute effect sizes higher than the minimum absolute effect size for significant null differential controls (marked by the dashed vertical line).

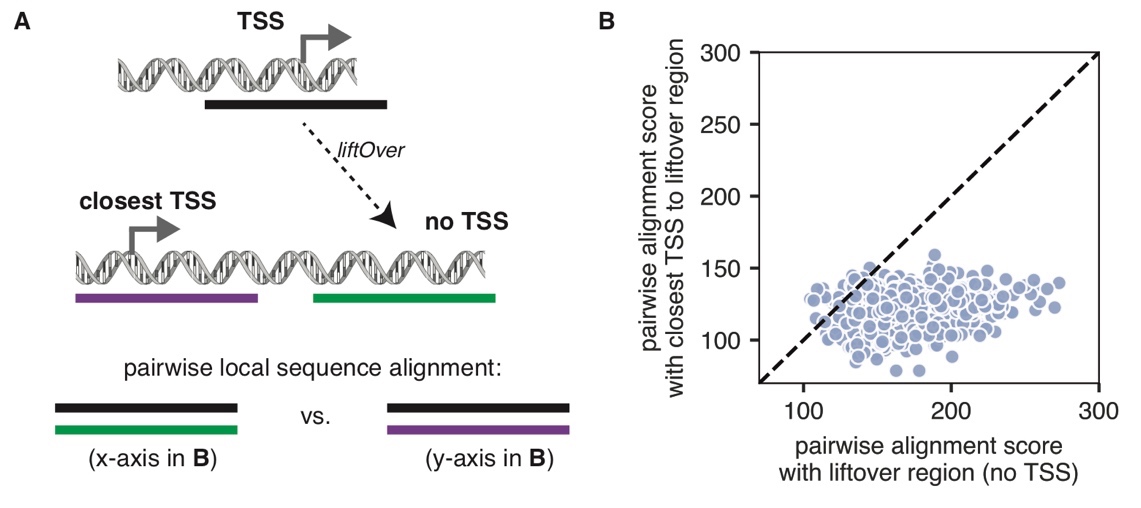

**Supplemental Figure S8: Pairwise local sequence alignment scores for non-conserved TSSs. A:** Schematic showing overview of pairwise alignments we computed: either the TSS to its orthologous region (as found via liftOver) that has no TSS, or the TSS to the closest TSS to its orthologous region (as found via liftOver). **B:** Scatter plot showing the sequence alignments outlined in A. Local sequence alignments were computed using the Biopython^10^ function localms from the pairwise2 module with the following parameters: identical characters = +2 points, non-identical character = -1 point, gap opening = -1 point, extending a gap = -0.1 points.

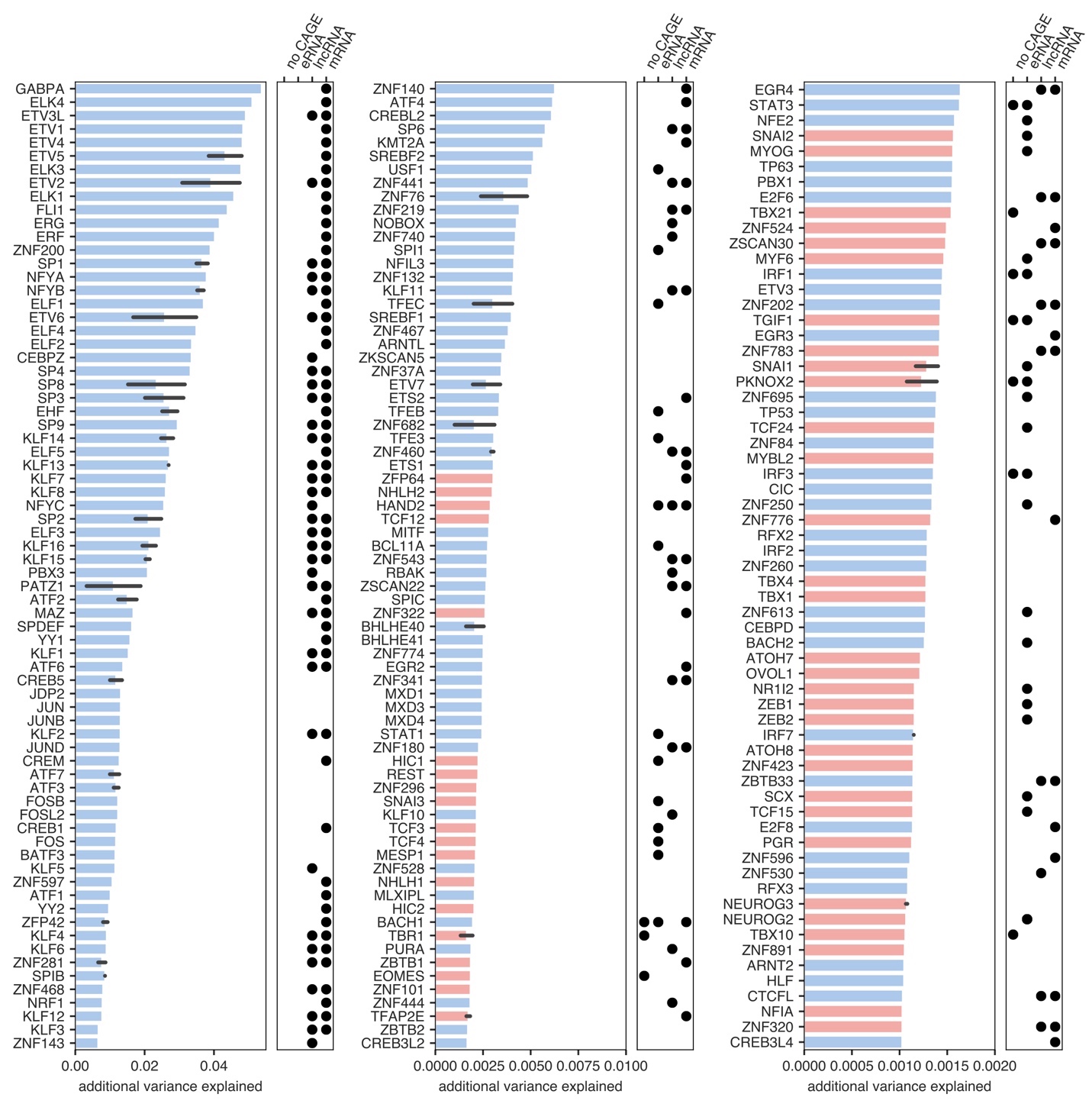

**Supplemental Figure S9: TF motifs significantly associated with MPRA activity.** Bar plots show the additional variance in mean MPRA activity explained by adding in a motif indicator variable. Color denotes whether motifs were predicted to be activators (blue) or repressors (red). When TFs had more than one “best” motif denoted in the Lambert et al catalog^11^, we plotted the average of all “best” motifs as well as its bootstrapped 95% confidence interval (gray bars). Dot plots show significant enrichment of motifs in a given biotype (adjusted p-value < 0.05), as denoted by a Hypergeometric test. In this plot, only the 241 significant motifs that explained ≥ 0.001 additional variance are shown. The full motif results can be found in **Supplemental Table S5**.

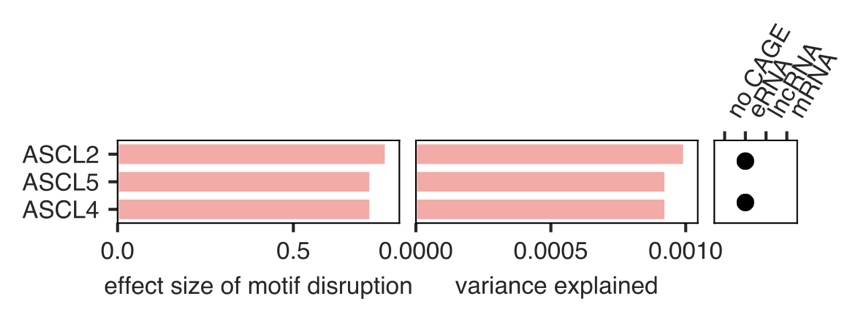

**Supplemental Figure S10: Repressive motifs significantly associated with *cis* effects.** Left: effect size associated with motif disruption. Right: enrichment of a given TF motif across biotypes, as determined by a Hypergeometric test. Black dots denote significant enrichment (FDR < 0.05).

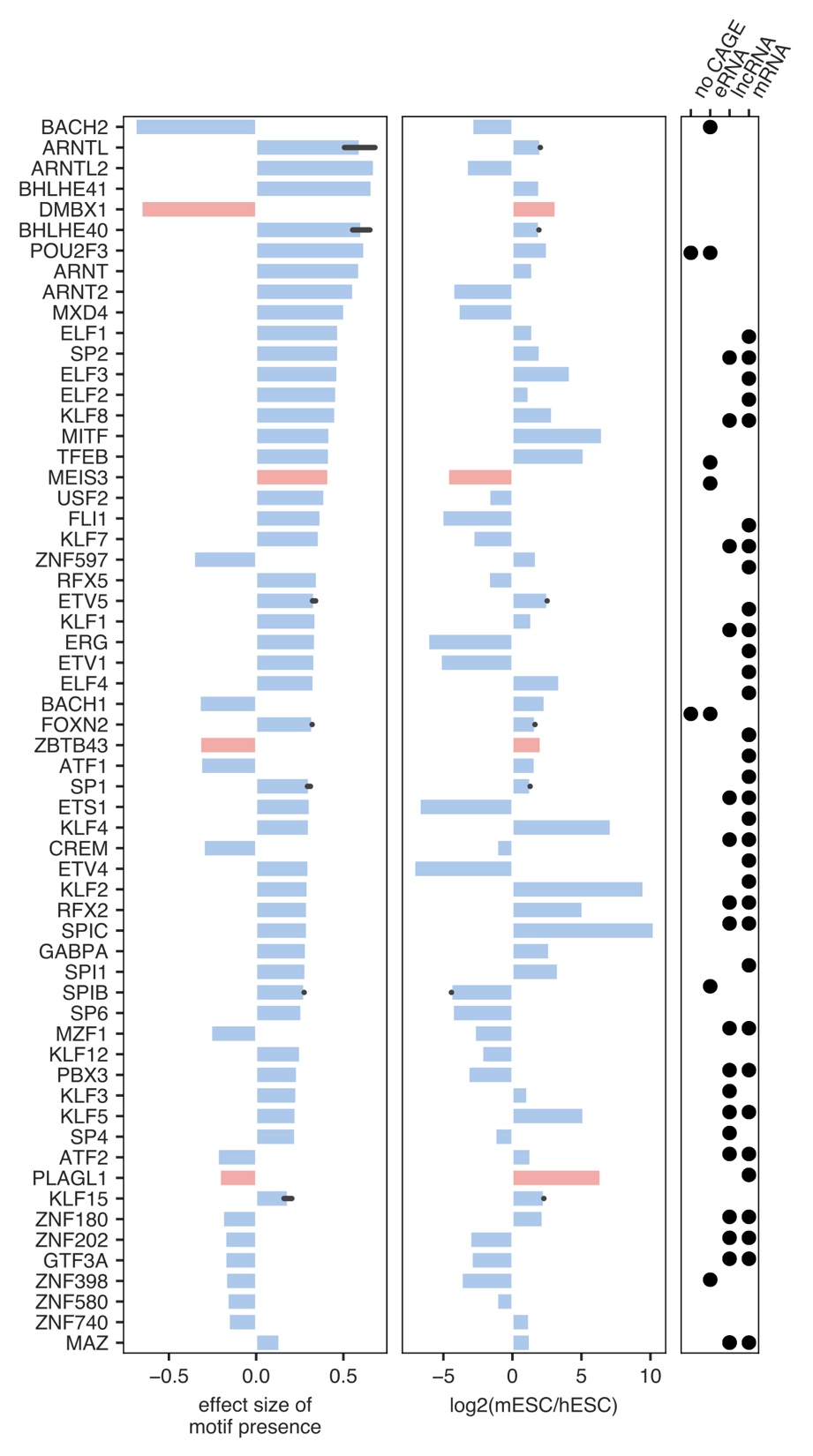

**Supplemental Figure S11: TF motifs significantly associated with *trans* effects that are also differentially expressed.** Plot showing the motifs significantly associated with *trans* effects (FDR < 0.05) that are also differentially expressed between hESCs and mESCs. Left: effect size associated with motif enrichment. Motifs that are associated with sequences more highly expressed in mESCs are > 0, and those associated with sequences more highly expressed in hESCs are < 0. Middle: log2 foldchange in expression via RNA-seq. Right: enrichment of a given TF motif across biotypes, as determined by a Hypergeometric test. Black dots denote significant enrichment (FDR < 0.05). Repressive motifs DMBX1, MEIS3, ZBTB43, and PLAGL1 all agree in their direction of enrichment in *trans* effects and their differential expression via RNA-seq.
